# Tension-induced suppression of allosteric conformational changes explains coordinated stepping of kinesin-1

**DOI:** 10.1101/2024.09.19.613825

**Authors:** Tsukasa Makino, Ryo Kanada, Teppei Mori, Ken-ichi Miyazono, Yuta Komori, Haruaki Yanagisawa, Shoji Takada, Masaru Tanokura, Masahide Kikkawa, Michio Tomishige

## Abstract

The dimeric motor protein kinesin-1 walks along microtubules by alternating ATP hydrolysis and movement of its two motor domains (“head”). The detached head preferentially binds to the forward tubulin-binding site after ATP binds to the microtubule-bound head, but the mechanism preventing premature binding to the microtubule while the partner head awaits ATP remains unknown. Here, we examined the role of the neck linker, the segment connecting the two heads, in this mechanism. High-resolution structural analyses of the nucleotide-free head revealed a bulge just ahead of the neck linker’s base that creates an asymmetric constraint on its mobility. While the neck linker can stretch freely backward, it must navigate around this bulge to extend forward. Based on this finding, we hypothesized that premature binding of the tethered head is suppressed by an intolerable increase in neck linker tension. Molecular dynamic simulations and single-molecule fluorescent assays supported this model. These findings demonstrate a tension-based regulation mechanism where off-pathway conformational transitions are thermodynamically suppressed through entropy loss associated with neck linker stretching, suggesting that neck linker tension influences the allosteric conformational transition rather than directly affecting the nucleotide state.

## Introduction

Kinesin-1 (hereafter referred to as kinesin) is a dimeric motor protein that transports cargo inside cells by moving toward the plus-end of the microtubule (Vale, 2003). Kinesin is a highly processive motor, capable of taking over 100 consecutive 8-nm steps before detaching from the microtubule. The unidirectional and processive motion of kinesin is powered by a hand-over-hand movement of two motor domains (“heads”) (Kaseda et al., 2003, Asbury et al., 2003; Yildiz et al., 2004). After the trailing head detaches from the microtubule, it surpasses the partner head and attaches to the forward-tubulin binding site located 16 nm ahead. The binding and dissociation of the head to and from the microtubule are tightly coupled to its ATP turnover; ADP release stabilizes the microtubule binding of the head, while ATP hydrolysis and phosphate release cause the head to detach from the microtubule (Cross, 2004; Cochran 2015). Consequently, for the kinesin to move hand-over-hand, the ATP hydrolysis cycles of the two heads must be out of phase.

In his elegant experiment, Hackney demonstrated that when a kinesin dimer encounters a microtubule, it releases ADP from only one of its heads (Hackney, 1994). Subsequent ATP binding to the microtubule-bound head is required to release ADP from the second head. This suggests that before ATP binding, one head is bound to the microtubule and is nucleotide-free, and the other head, still bound to ADP, is tethered by its partner. ATP binding to the microtubule-bound head then aids the tethered head in attaching to the microtubule.

To illustrate the ATP-induced stepping of the tethered head, a model involving a conformational change in the neck linker region, a 14 amino-acid stretch that connects the two heads, has been proposed (Vale and Milligan, 2000). The neck linker has been shown to undergo a disordered-to-ordered structural transition depending on the nucleotide state of the head (Rice et al., 1999). In the ATP or ADP·Pi state, the neck linker is docked onto the head, with its distal end pointing forward, whereas in the nucleotide-free or ADP state, it detaches from the head, becoming disordered. This discovery led to the model that docking of the neck linker to the microtubule-bound head upon ATP-binding moves the tethered head forward, and the subsequent diffusional search of the tethered head allows it to bind to the forward-tubulin binding site. This mechanism explains how the tethered head preferentially binds to the forward-binding site after ATP binding to the microtubule-bound, leading head. However, it does not clarify how the tethered head is prevented from rebinding to the rear-tubulin binding site before ATP binds to the leading head.

The original neck linker docking model, therefore, hypothesized that the ADP-bound trailing head, after Pi release, is not away from the tubulin-binding site; instead, it’s still weakly bound to the microtubule, maintaining a two-head-bound state (Hancock and Howard, 1999; Vale and Milligan, 2000; Shief et al., 2004). The neck linker docking of the leading head facilitates the dissociation of the trailing head from the microtubule, translating the head forward. However, three laboratories using different single-molecule techniques led to the conclusion that under limited ATP concentrations, the kinesin dimer stably assumes a one-head-bound state, in which the trailing head is displaced from the rear-tubulin binding site while waiting for ATP binding (Mori et al., 2007; Guydosh and Block, 2009; Asenjo et al., 2009). More recent studies with high temporal resolution single-molecule observations of one of the kinesin heads further demonstrated that kinesin waits for ATP-binding in the one-head-bound state under a wide range of ATP concentrations, including saturating ATP conditions (Isojima et al., 2016; Wolff et al., 2023).

These experiments provided in-depth information about the structural state of the ATP waiting state; the tethered head is positioned on average to the right and slightly behind the microtubule-bound head (Mori et al., 2007; Isojima et al., 2016; Wolff et al., 2023), and has a significant translational and rotational freedom (Guydosh and Block 2009; Asenjo et al., 2009; Isojima et al, 2016). These findings about the ATP-waiting state provoke a question: How is the tethered head’s rebinding to the rear-tubulin binding site prevented before ATP-binding to the microtubule-bound head? When the neck linkers were artificially extended by inserting spacers between the neck linker and the coiled-coil stalk, the kinesin dimer could release both ADPs rapidly, even without added ATP (Hackney et al., 2003; Andreasson et al., 2015), and the tethered head could bind to rear and forward-tubulin binding sites before ATP-binding to the microtubule-bound head (Yildiz et al., 2008; Isojima et al., 2016). In the two-head-bound state of wild-type kinesin, the neck linkers of the front and rear heads are stretched in opposite directions due to their limited length (Benoit et al., 2021). These findings suggest that constraints on the neck linker (whether from steric hindrance or interactions with the head or microtubule) are crucial in preventing the tethered head from binding to microtubule.

To understand the mechanism that prevents premature binding of the tethered head to the microtubule during ATP-waiting state, we first studied the neck linker conformation in the nucleotide-free state using X-ray crystallography and cryo-electron microscopy (cryo-EM). We found that the motility of the neck linker in the nucleotide-free state is asymmetrically constrained due to a steric hindrance from the motor domain, which impacts the neck linker’s extensibility toward the plus-end of the microtubule. Using this structural information, we built a model of dimeric kinesin in a two-head-bound state where both heads are nucleotide-free. Comparing this with the two-head-bound state with an ATP-bound trailing head and a nucleotide-free leading head, we theorized that excessive increase in neck linker tension prevents the transition to the two-head-bound state where both heads are nucleotide-free. We tested this hypothesis using molecular dynamic (MD) simulation and single-molecule fluorescence resonance energy transfer (smFRET). These results suggest a tension-based mechanism that controls the transition from one-head-bound to two-head-bound states, as well as transitions between two-head-bound states; among possible conformational transitions, the one that requires less entropy reduction from stretching the disordered neck linker is favored.

## Results

### Structural analysis of the neck linker in nucleotide-free state

The crystal structure of the nucleotide-free kinesin-1 motor domain has been solved as a complex with tubulin (Cao et al., 2014) and as mutant kinesins (Cao et al., 2017). However, the neck linker was truncated in these constructs, leaving the high-resolution structure of the kinesin-1 neck linker in the nucleotide-free state unclear. Here, we analyzed the structure of a human kinesin-1 construct of 336 amino acids, which includes the N-terminal motor domain (1-322 residues) and the subsequent neck linker (323-336 residues), using X-ray crystallography. For consistency across experimental techniques and comparison with the previously solved nucleotide-free kinesin-1 structures, we used a cysteine-light mutant kinesin, where surface-exposed cysteines were replaced with either Ala or Ser (Rice et al., 1999; Tomishige and Vale, 2000). Since kinesin in solution tightly binds ADP and becomes unstable in the absence of both nucleotide and microtubule (Cross, 2004), we used a G234A mutant kinesin, in which Gly234 residue in the switch II loop was replaced with Ala. This mutant is known to rapidly release ADP, even in the absence of microtubules (Rice et al., 1999).

First, we verified that the G234A and cysteine-light mutations did not alter the structure of the kinesin motor domain in an ADP-bound state (Fig. 1 A). We then successfully crystallized the G234A mutant kinesin under a nucleotide-free condition. The X-ray crystal structure was determined at a 2.8 Å resolution (Fig. 1 B; and Table 1). The electron density map of the nucleotide-binding pocket revealed the absence of a nucleotide or an ion that mimics inorganic phosphate (Fig. S1). The overall structure of the G234A motor domain was similar to that of the previously solved nucleotide-free kinesin-1 (Cao et al., 2014) (Fig. 1 B). The neck linker, directly following the α6-helix of the motor domain, was only visible for the initial two residues (K323 and T324) (Fig. 1 C); the rest of the neck linker was disordered, consistent with the previous reports (Rice et al., 1999; Tomishige et al., 2006). The observed neck linker structure was independent of crystal packing effects, as the nearest element of the adjacent chain (L5) was 1.1 nm away from the K324 residue. The position of T324 is stabilized by a hydrogen bond with the side chain of R321 residue of α6. In the ADP·AlF4-bound state, these residues fold into an extra turn of α6-helix (Fig. 1 D), while in the nucleotide-free state, the α4 helix, which lies vertically to the α6 helix, sterically prevents the α-helix formation of these residues. Consequently, the unfolded K323 and T324 residues are excluded from the motor domain and point away from the core (Fig. 1 C). Similar results were obtained using a crystal structure of G234V mutant (Fig. S2).

**Figure 1.**
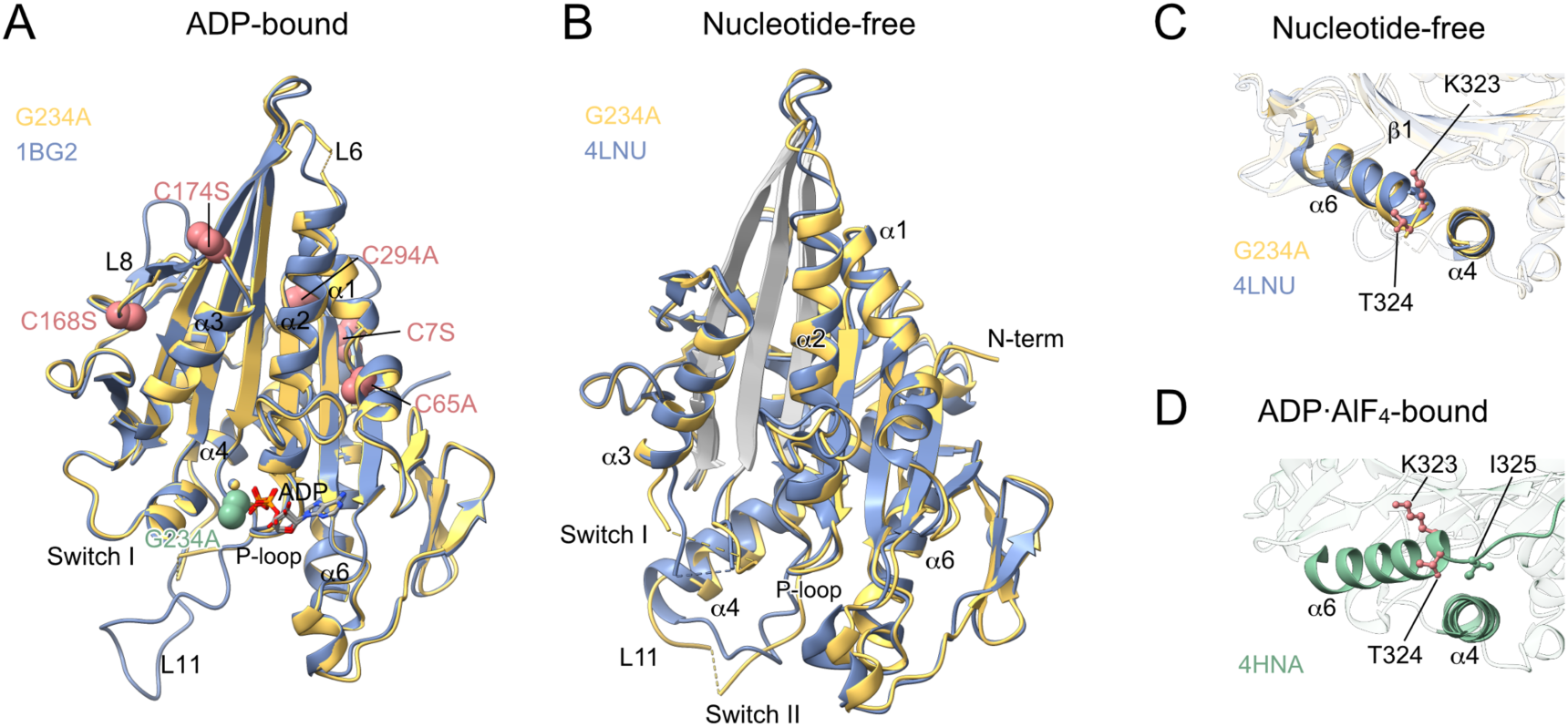
Crystal structures of G234A mutant kinesin-1 motor domain. **(A)** 2.4 Å crystal structure of Cys-light mutant kinesin-1 (336 amino-acids) with G234A mutation complexed with ADP (yellow), aligned with the ADP-bound structure of wild-type human kinesin-1 (blue; PDB# 1BG2 (Kull et al., 1996)). The root mean square deviations (RMSD) of backbone atoms between these structures was 0.819 Å. The G234A residue is highlighted as a green sphere, and cysteine substitutions are shown as red spheres. **(B)** 2.8 Å crystal structure of nucleotide-free G234A mutant kinesin-1 motor domain (yellow), aligned with the nucleotide-free motor domain from the kinesin-tubulin complex crystal structure (blue; #4LNU (Cao et al., 2014)). The RMSD of backbone atoms between these structures was 1.259 Å. L11 is disordered in our crystal structure but ordered in 4LNU, suggesting that microtubule binding is necessary for L11 folding. **(C and D)** Close-up views of the α4 and α6 helices in nucleotide-free and ADP·AlF_4_-bound (green; #4HNA (Gigant et al., 2013)) states. The initial residues of the neck linker (K323, T324, and I325) are highlighted as ball and stick models.

**Table 1.** Data collection and refinement statistics of the crystal structures. *Nucleotide-free G234A and G234V crystal contains two chains in an asymmetric unit. ^#^Numbers in parenthesis show resolution in the highest resolution shell.

|  | G234A<br>(nucleotide-free) | G234A<br>(ADP) | G234V<br>(nucleotide-free) |
| --- | --- | --- | --- |
| **Data collection** |  |  |  |
| Space group | $P2_1$ | $P2_1$ | $P2_1$ |
| Cell dimensions |  |  |  |
| $a, b, c$ (Å) | 44.2, 156.9, 53.0* | 49.4, 67.9, 55.5 | 44.1, 162.4, 53.2* |
| $\beta$ (°) | 114.6 | 96.1 | 114.5 |
| Resolution (Å) | 20-2.82 (2.82-2.90)<sup>#</sup> | 20-2.40 (2.40-2.46) | 20-2.80 (2.87-2.80) |
| $R_{\text{merge}}$ | 0.062 (0.098) | 0.052 (0.412) | 0.051 (0.076) |
| $I/\sigma I$ | 16.13 (4.96) | 17.90 (3.73) | 29.58 (13.25) |
| Completeness (%) | 95.5 (57.5) | 99.7 (100) | 97.6 (85.6) |
| Redundancy | 3.4 (1.3) | 4.2 (4.2) | 7.1 (4.6) |
| **Refinement** |  |  |  |
| Resolution (Å) | 19.6-2.82 | 20-2.40 | 19.5-2.80 |
| No. reflections | 15129 | 14330 | 16383 |
| $R_{\text{work}}/R_{\text{free}}$ (%) | 20.7/ 26.5 | 19.2/ 25.2 | 20.0/ 25.8 |
| No. atoms |  |  |  |
| Protein | 4761 | 2363 | 5017 |
| ADP/ion | 0/0 | 27/1 | 0/0 |
| Water | 1 | 53 | 0 |
| $B$-factors (Å<sup>2</sup>) | | | |
| Protein | 24.2 | 45.0 | 27.5 |
| ADP/ion |  | 33.6/27.4 |  |
| Water | 12.1 | 45.0 |  |
| R.m.s deviations |  |  |  |
| Bond lengths (Å) | 0.004 | 0.019 | 0.003 |
| Bond angles (°) | 0.61 | 1.86 | 0.54 |

Next, we analyzed the structure of kinesin in complex with microtubule using cryo-EM to elucidate how the microtubule influences the neck linker configuration. Cysteine-light kinesin-1 construct of 349 amino acids was bound to microtubule under nucleotide-free conditions, and its structure was reconstituted at a 3.48 Å resolution (Fig. 2 A-C; Fig. S3), better than that of the previously reported cryo-EM structure of nucleotide-free kinesin-1 (Shang et al., 2014). The atomic model was built using a flexible fitting with the crystal structures of the nucleotide-free G234A and GMPPNP-bound microtubule. The refined model closely resembles the previously reported crystal structure of nucleotide-free kinesin-tubulin complex (Cao et al., 2014), except that the N-terminus of α4, which extends upon microtubule-binding, along with L11, is nearly straight in our structure but distorted toward the plus-end in the Cao et al.’s structure (Fig. 2, D and E). The discrepancy can be explained by the fact that Cao et al.’s kinesin is bound to curved tubulin, whereas our cryo-EM model is on the straight microtubule and better describes the interactions in the kinesin-tubulin interface (Fig. 2 F-H). The neck linker is mostly disordered, with only the first two residues, K323 and T324, visible (Fig. 2, B and C), consistent with results from the G234A mutant’s crystal structure. These findings suggest that the neck linker doesn’t interact specifically or stably with either the motor domain or the microtubule surface in the nucleotide-free state.

**Figure 2.**
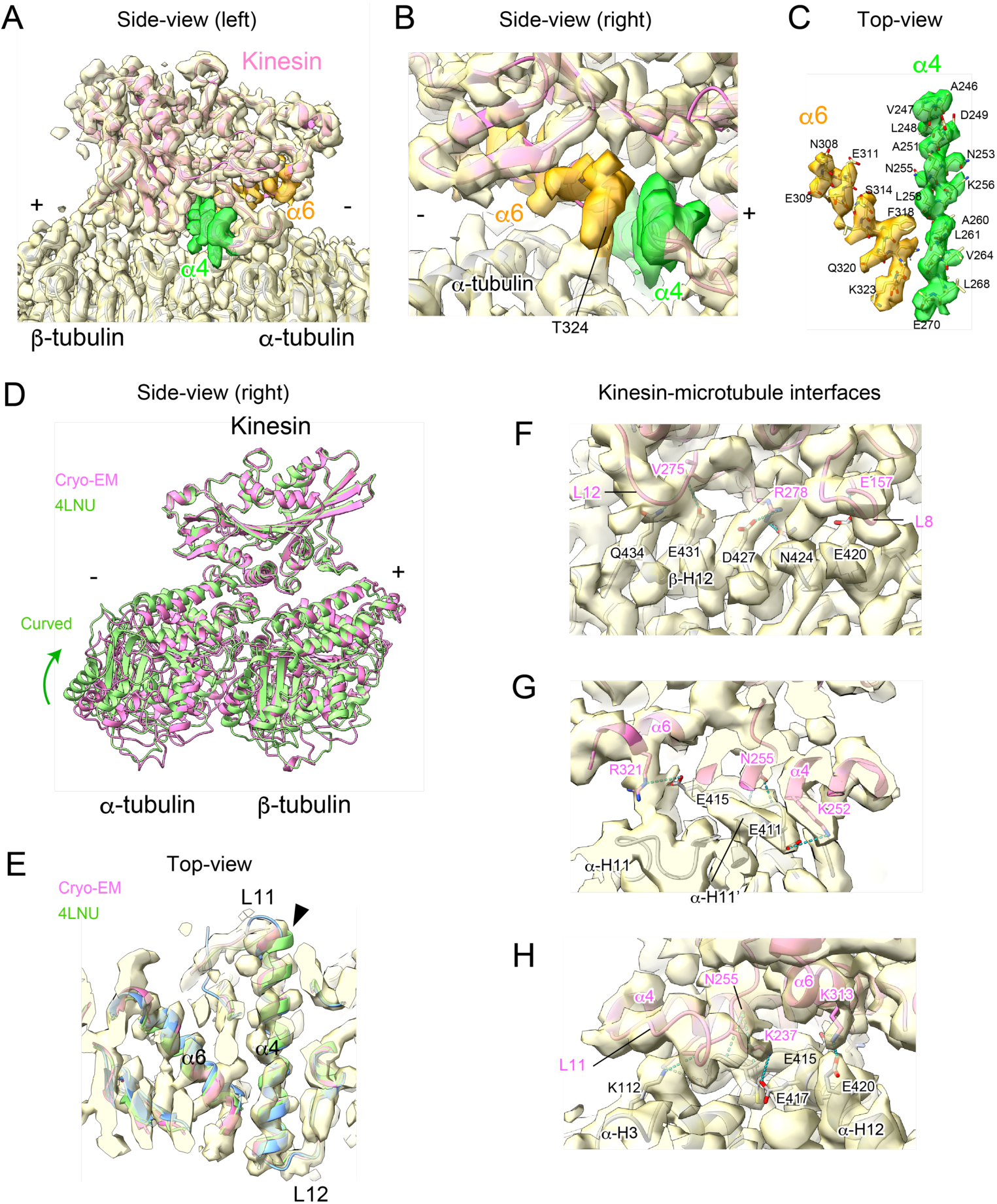
Cryo-EM density map and atomic model of nucleotide-free kinesin on the microtubule. **(A-C)** Cryo-EM reconstructed map of the nucleotide-free kinesin-microtubule complex (light yellow) and its refined atomic model (kinesin; pink, tubulin; gray) are shown in cartoon representation (A and B) and stick representation (C). The cryo-EM map of α4 and α6 helices are highlighted in green and orange, respectively. Panels B and C provide close-up views of α4 and α6. (**D and E**) The atomic model derived from the cryo-EM map was aligned with the previously solved nucleotide-free structure of human kinesin-1 motor domain bound to tubulin-DARPin complex (green; #4LNU (Cao et al., 2014)). These structures were similar except that the N-terminus of α4 of 4LNU deviated from the cryo-EM density map, as shown by the arrowhead in panel E. **(F-H)** Close-up views of the cryo-EM map and its atomic model of the kinesin-microtubule interface. The interface of kinesin with β-tubulin (F) and α-tubulin (G and H) is shown. Hydrogen bonds and salt bridges are marked as blue dashed lines.

### Structural feature of nucleotide-free motor domain peripheral to the neck linker

We then turned our attention to the structural feature of the kinesin motor domain unique in the nucleotide-free state, specifically those parts adjacent to the neck linker, and studied how these parts influence the configuration of the disordered neck linker. In previous research, Grant et al. classified 114 distinct chains from 69 kinesin family crystal structures using principal component analysis and found that they are segregated into two major groups (termed ATP-like and ADP-like conformational states) along with a minor cluster that includes exclusively inhibitor-bound kinesin-5 (Grant et al., 2007; Scarabelli et al., 2013). Here, we performed the principal component analysis on 247 polypeptide chains from 137 kinesin family crystal structures deposited in the Protein Data Bank, which includes the nucleotide-free crystal structures by Cao et al., plus our nucleotide-free structures of kinesin-1 (G234A and G234V crystal and wild-type cryo-EM structures). The 2D conformer plot of the first two principal components shows that our nucleotide-free structures and those of Cao et al. form a distinct cluster separate from the ATP-like and ADP-like states (Fig. 3, A and B), demonstrating that these nucleotide-free structures represent a third conformational state of kinesin structure. Other crystal structures of kinesin family proteins without bound nucleotide (but with an ion bound to P-loop that may mimic phosphate) are classified into ADP-like or ATP-like states (Shipley et al., 2004; Morikawa et al., 2015; Guan et al., 2017; Cao et al., 2017).

**Figure 3.**
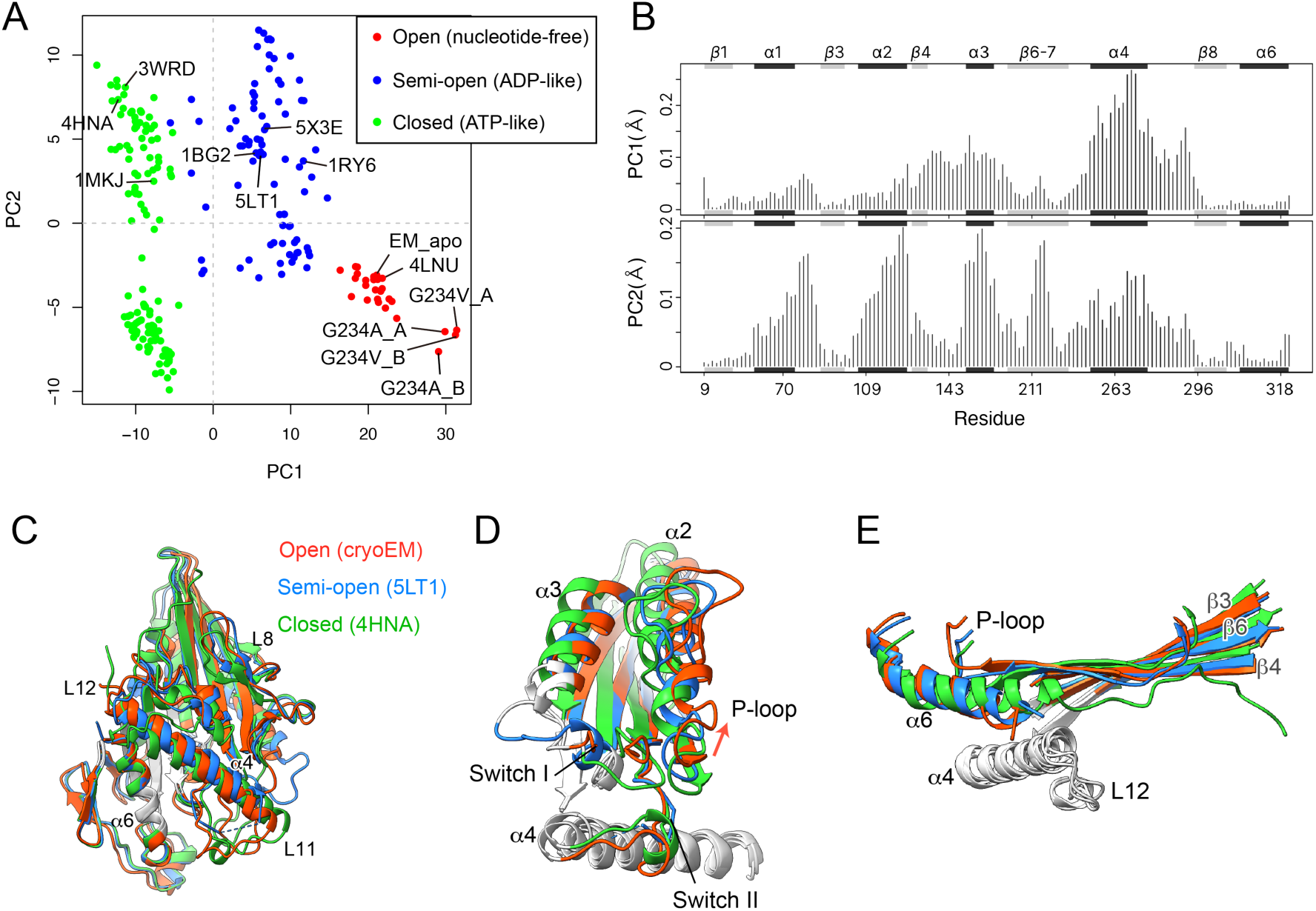
Structural classification of kinesin motor domain using PCA. **(A)** Principal component analysis (PCA) classified kinesin crystal structures based on their conformational similarity. We projected kinesin structures from the Protein Data Bank—including our crystal and cryo-EM apo structure—onto principal planes defined by the two most significant principal components (PC1 and PC2). Each dot’s color represents one of three classes obtained from hierarchical clustering of the projected structures in the PC1 to PC4 planes. Representative structures of kinesin-1 for each class (1BG2, 1MKJ, 4HNA, 4LNU) and previously reported nucleotide-free structures (1RY6, 3WRD, 5LT1, 5X3E), are labeled with their PDB code. **(B)** The contribution of each residue (as for human kinesin-1) to the first two principal components (PC1 and PC2). Shaded rectangles indicate the corresponding secondary structures. **(C-E)** Comparison of representative structures from three classes. 5LT1 was chosen to represent the semi-open state, as the N-terminus of its α4 helix extends similarly to the microtubule-bound states (cryoEM apo and 4HNA). These three structures are superposed on the invariant “core” (gray in C) or the B-domain (gray in D and E). These structures were used to identify subdomains (see Fig. 4).

The correlation between the bound-nucleotide and the conformational state of kinesin crystal structures is not evident in the absence of microtubule (e.g., the ATP-like class primarily consists of ADP-bound crystal structures). Moreover, a recent cryo-EM analysis of KIF14 on the microtubule revealed that the AMP-PNP-bound motor domain could adopt a conformation similar to the nucleotide-free head when its neck linker is truncated or detached from the head (Benoit et al., 2021). Therefore, we adopted the nomenclature of kinesin’s three conformational states proposed by Sosa’s group, which is based on the openness of the nucleotide-binding pocket rather than the bound-nucleotide (Benoit et al., 2023); we termed these three classes as open (which includes our nucleotide-free structures), semi-open (formally known as ADP-like), and closed (previously known as ATP-like) states (Fig. 3, C-E). We chose the term “semi-open” over “semi-closed” to describe the ADP-like state because the angle of the subdomain, which influences the openness of the nucleotide pocket, in the semi-open state is rather similar to that of the open state (will be described later).

The conformational changes from open to closed states in kinesin-1 can be approximated as relative rigid-body movements of three distinct subdomains, as demonstrated by Cao et al. (Cao et al., 2014), although this previous analysis did not include the semi-open state. Here, we identified the subdomains of kinesin-1 by comparing representative structures from the open, closed, and semi-open states using a domain-search program (Fig. 4 A). The three subdomains we identified differ from those identified by Cao et al.; one of the domains remains nearly stationary on the microtubule during the conformational changes between three states (we refer to this as Bottom-domain or B-domain), and the other two domains move relative to the stationary B-domain during the conformational changes (Fig. 4 B), which are spatially divided into front and rear halves of the motor domain, with the central α4 helix as a border (Fig. 4, C and D); therefore, we termed Front- and Rear-domains, or F- and R-domains.

**Figure 4.**
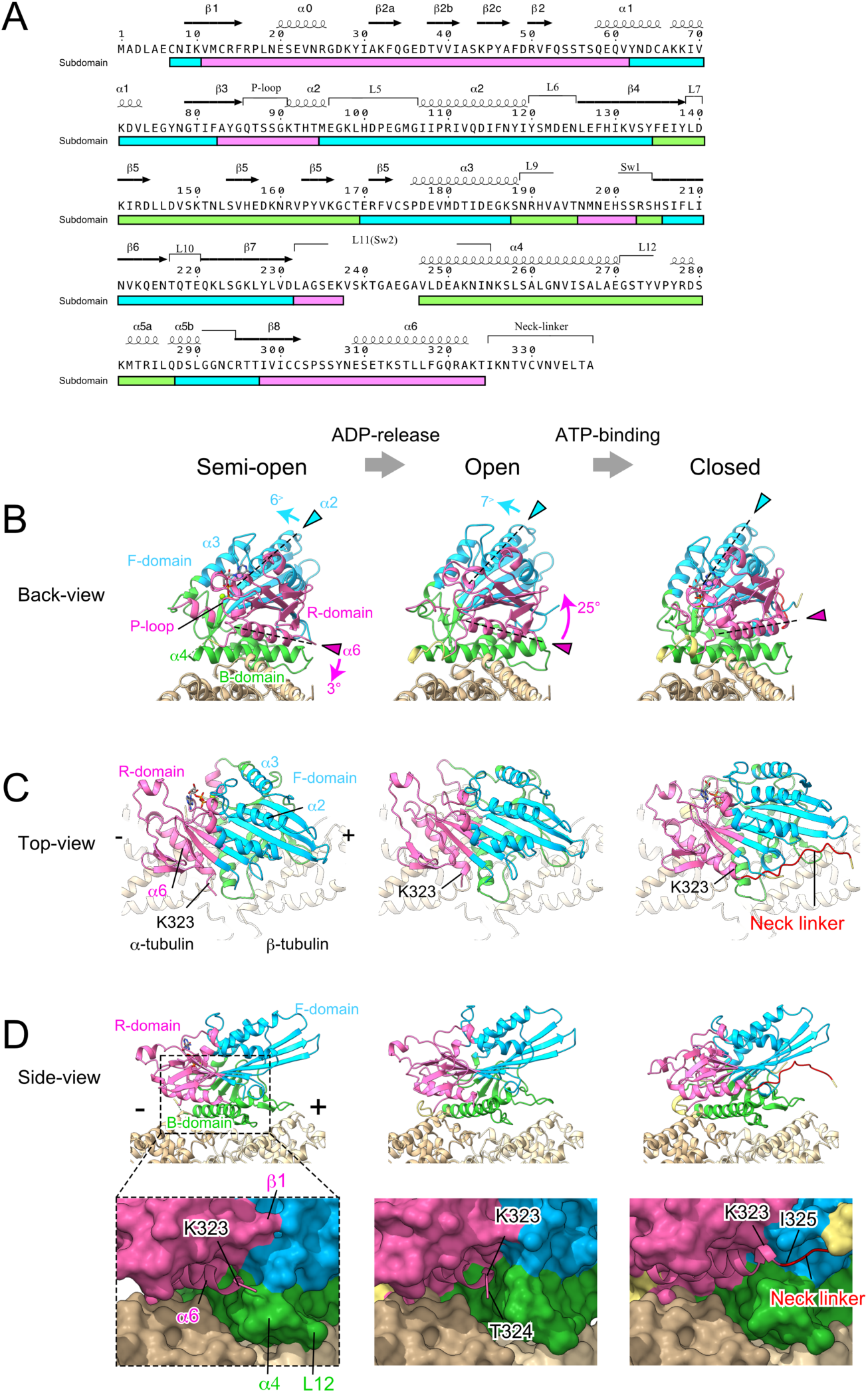
Subdomain motions associated with the allosteric conformational changes of kinesin motor domain on the microtubule. **(A)** The residues classified into three subdomains are colored in magenta (R-domain), cyan (F-domain) and green (B-domain). **(B-D)** Three classes of conformational state of the kinesin motor domain (Fig. 3) bound to the microtubule, colored by subdomain. Unassigned residues, which are disordered in the semi-open state, are yellow and the neck linker is red. The kinesin-microtubule complex models of semi-open and closed states were prepared by fitting 5LT1 and 4HNA onto the cryo-EM density maps of ADP-bound (Sindelar and Downing, 2010) and ADP·AlF_x_-bound (Shang et al., 2014) states, respectively. For clarity, we depicted the semi-open state with MgADP bound by incorporating the MgADP from the 1BG2 structure into the 5LT1 (nucleotide-free) structure. The rotational movements of R- and F-domains relative to the microtubule—from semi-open to open and from open to closed—are represented by angular changes in the long axes of α6 and α2 helices (arrowheads), respectively.

### Nucleotide-dependent subdomain motions affect the neck linker mobility

The transition from semi-open to open conformation, coupled with the release of ADP from the motor domain after microtubule-binding, results in anticlockwise rotational movements of the F-domain when viewed from the minus-end, while the R-domain rotates clockwise (Sindelar, 2011; Cao et al., 2014; Benoit et al., 2023; Fig. 4B; Video 1). The rotational movement of the R-domain (containing P-loop and α6 helix; Fig. 4 A) relative to the B-domain (including α4 helix) leads to the opening of the nucleotide-binding pocket located on the left side, and a steric clash between the C-terminus of α4 on the right side and the distal end of α6 where the neck linker is linked (Fig. 4, B and C). This steric hindrance likely imposes an asymmetric constraint on the mobility of the disordered neck linker (Fig. 4 D; Video 2). Specifically, the neck linker can stretch freely backward (toward the microtubule’s minus-end), while its forward mobility is restricted by the steric hindrance from the C-terminus of α4 helix and subsequent L12.

To explore the hypothesis that the neck linker’s mobility is asymmetrically constrained in the nucleotide-free/open state, we visualized the position of the mobile neck linker in the nucleotide-free state by labeling it with a gold cluster (Rice et al., 1999; Kikkawa et al., 2000). To comprehend the influence of steric hindrance near the neck linker’s proximal end, a maleimide-modified gold cluster with a diameter of 1.4 nm was attached to the thiol residue of T328C in a cysteine-light kinesin head, which is four amino acids away from the last ordered neck linker’s residue T324. A cryo-EM density map with a resolution of 15 Å showed that the density corresponding to the gold cluster appeared as an ellipsoidal shape extending along the microtubule protofilament (Fig. 5 A). The centroid of the gold density is 2.37 nm away from the T324 residue, a distance consistent with the combined length of a 4 amino-acids peptide (approximately 1.5 nm), the maleimide linker, and the gold cluster’s radius. The gold density is located rearward from the proximal end of the neck linker or the distal end of α6 helix and away from the microtubule surface. These observations support the hypothesis that, in the absence of a bound nucleotide, the mobility of the disordered neck linker is biased toward the minus end of the microtubule. This finding is also consistent with the right and rearward positioning of the tethered head relative to the microtubule-bound head in kinesin dimer during the ATP-waiting (Fig. 5 B; Mori et al., 2007; Isojima et al., 2016; Wolff et al., 2023).

**Figure 5.**
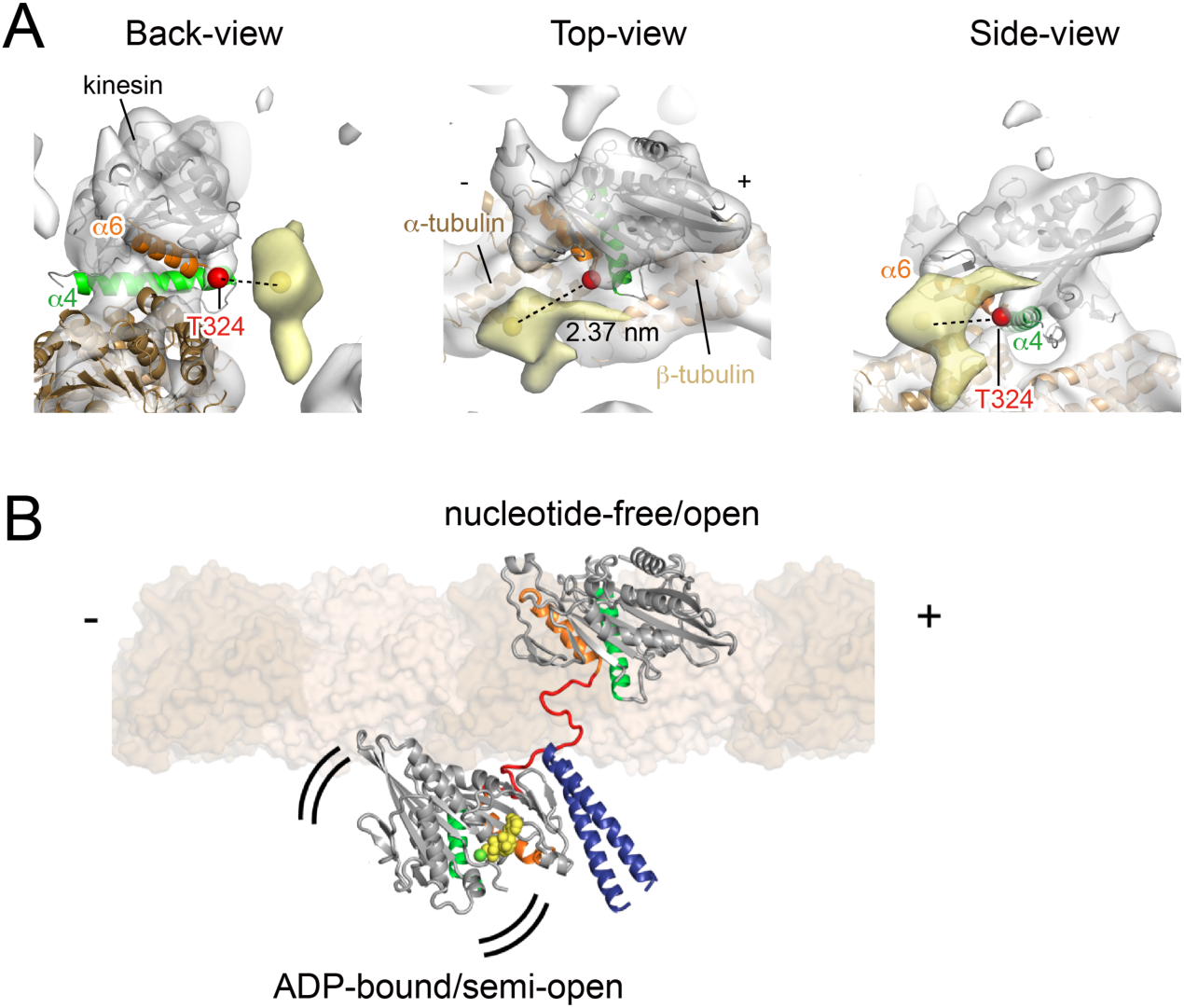
Cryo-EM observation of the proximal end of the neck linker in the nucleotide-free state. **(A)** 15 Å cryo-EM density map (σ =1.0) of gold-labeled kinesin-microtubule complex under nucleotide-free condition. A gold cluster was specifically attached to the Cys328 residue on the neck linker. α4 and α6 helices are shown in green and orange, respectively. Red spheres indicate the last ordered residue of the neck linker in the nucleotide-free state (T324). Nucleotide-free kinesin and tubulin crystal structures have been fit to the density map. Extra density corresponding to the attached gold is highlighted in light yellow. The centroid of the gold cluster, identified as the maximum density region (σ =2.7), is shown as yellow spheres. **(B) S**tructural model of kinesin dimer in the one-head-bound state: the microtubule-bound head is nucleotide-free and in the open state, while the unbound head is in the semi-open state, tethered by disordered neck linkers. The neck linker and the neck coiled-coil are shown in red and blue, respectively.

### Structural basis to prevent rebinding of the tethered head while waiting for ATP

The asymmetric constraint on the neck linker in the nucleotide-free/open state provides insights into the mechanism that prevents the transition to the two-head-bound state where both heads are nucleotide-free and in open conformation (termed Topen-Lopen state) while waiting for ATP binding. Given that the contour length of one of the unfolded neck linkers is about 5.3 nm (assuming a contour length of 3.8A per amino acid (Hyeon and Onuchic, 2007; Hariharan and Hancock, 2009)), when both heads of kinesin dimer bind to adjacent tubulin-binding sites separated by 8.2 nm, the neck linker in the leading head stretches backward, while the one in the trailing head stretches forward (Benoit et al., 2021). We created a model of the kinesin dimer on a microtubule in the Topen-Lopen state, where both heads are nucleotide-free and adopt the open conformation. First, the atomic models of two nucleotide-free heads were placed on the adjacent tubulin-binding sites and were connected by disordered neck linker chains (Fig. S4, A and B). Then, a typical configuration of the disordered neck linker chains in the thermal equilibrium state was estimated using MD simulations (Fig. S4 C; Video 3).

The neck linker of the leading nucleotide-free/open head stretches backward without steric hindrance; however, that of the trailing open head cannot go straight toward the plus-end of microtubule due to the steric hindrance posed by the α4 helix (Fig. 6 A, left). In the equilibrium configuration, the neck linker in the trailing head detours around to extend forward. Throughout the simulation, the stretched neck linker remained displaced from the head without any interaction, suggesting that the neck linker behaves as an entropic spring. To illustrate the difference between the neck linkers of trailing head in the open and closed conformational states, we also modeled the kinesin dimer in the two-head-bound state where the trailing head is ATP-bound and in the closed conformation (Tclose-Lopen; Fig. 6 A, right; Video 4). During this simulation, we observed a stable contact between the K326 side chain of the disordered neck linker and the D37 and F48 residues of the leading head (see Video 4), suggesting that the neck linker tension in Tclose-Lopen state includes an energetic component. The distance between T328 residue in the trailing head and I325 residue in the leading head increases by 0.96 nm in Topen-Lopen compared to Tclosed-Lopen (Fig. 6 A). Given that the disordered polypeptide chain acts like an entropic spring, extending the end-to-end distance of the disordered neck linker increases the tension applied to both ends. To estimate the amount of this tension, we isolated the disordered neck linker segments from both leading and trailing heads that stretch between the motor domains without steric hindrance or docking onto the head (Fig. S4 D). Then, we applied a harmonic potential to the Cα atoms at both ends of the stretched region and calculated the tension from the average displacement of the Cα atom from the potential minimum using MD simulations (Fig. 7, A and B). The calculated tension on the neck linker in Topen-Lopen was 98 pN, which is twice as much as in the Tclosed-Lopen state (39 pN; Fig. 7C). The tension in the Tclosed-Lopen state is likely an overestimate since this measurement excludes the enthalpic component discussed above, though it is comparable to previous MD measurements and theoretical calculations using a worm-like chain model (Hariharan and Hancock, 2009). Therefore, we hypothesized that the transition from the one-head-bound state to the Topen-Lopen two-head-bound state is thermodynamically unfavorable because it would result in an intolerable increase in the tension applied to the neck linker, while the transition to the Tclosed-Lopen state is feasible, as the tension applied to the neck linker is lower and might be tolerable.

**Figure 6.**
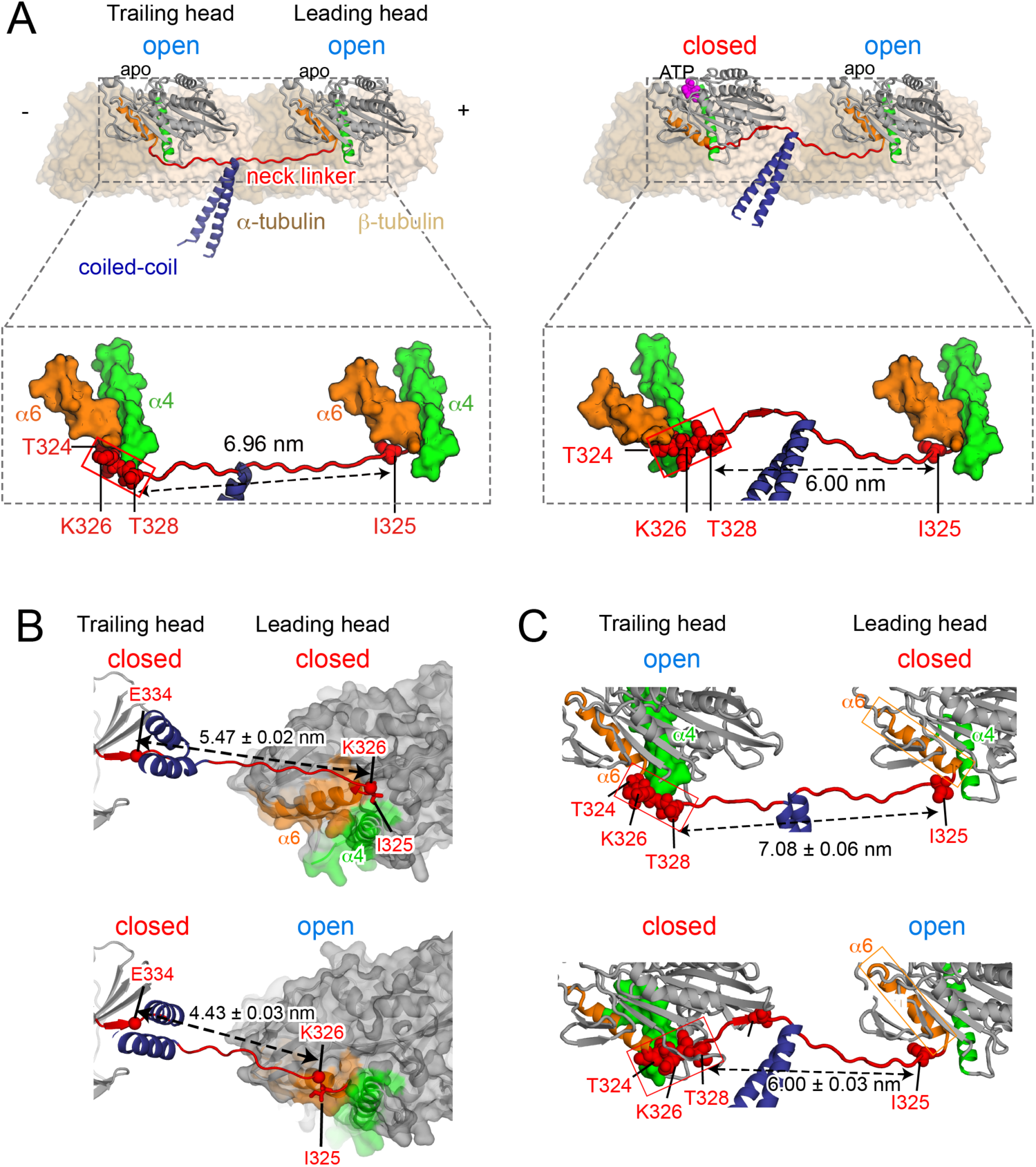
Structural models of dimeric kinesin on microtubule in all four combinations of open and closed heads. **(A)** Atomic models of kinesin dimer on microtubule in the two-head-bound state (top view): one with both heads nucleotide-free and in open state (left; termed T_open_-L_open_) and another with the trailing head ATP-bound and in closed state, while the leading head remains the nucleotide-free/open state (right; termed T_closed_-L_open_). Thermally equilibrium conformations of the disordered neck linkers were determined by running MD simulations (Fig. S4). The lower panel shows close-ups of the neck linkers (red), α4 (green) and α6 (orange) helices. Numbers indicate the distance between the T328 residue in the trailing head and the I325 residue in the leading head. (**B**) Close-up view (side view) of the atomic model of two-head-bound state with both heads in closed state (upper; T_closed_-L_closed_), compared with the T_closed_-L_open_ model (lower). We modeled the T_closed_-L_closed_ such that the initial segment of the neck linker (323-325 residues) of the leading head docks onto the head, while the rest remains detached. Even this partial docking leads to a considerable increase in neck linker extension, prohibiting the open-to-closed conformational change of the leading head (Fig. 9, purple rectangle). **(C)** Atomic model of T_open_-L_closed_ state (upper), compared with the T_closed_-L_open_ model (lower). This model represents the off-pathway transition where the tethered head binds to the rear-tubulin binding site after ATP-induced isomerization of the microtubule bound head (Fig. 9, green hexagonal box). In this model, the I325 residue of the leading head is detached from the head, but its position is translated toward the plus-end of microtubule due to the rotational movement and extension of the α6 helix (orange rectangle).

**Figure 7.**
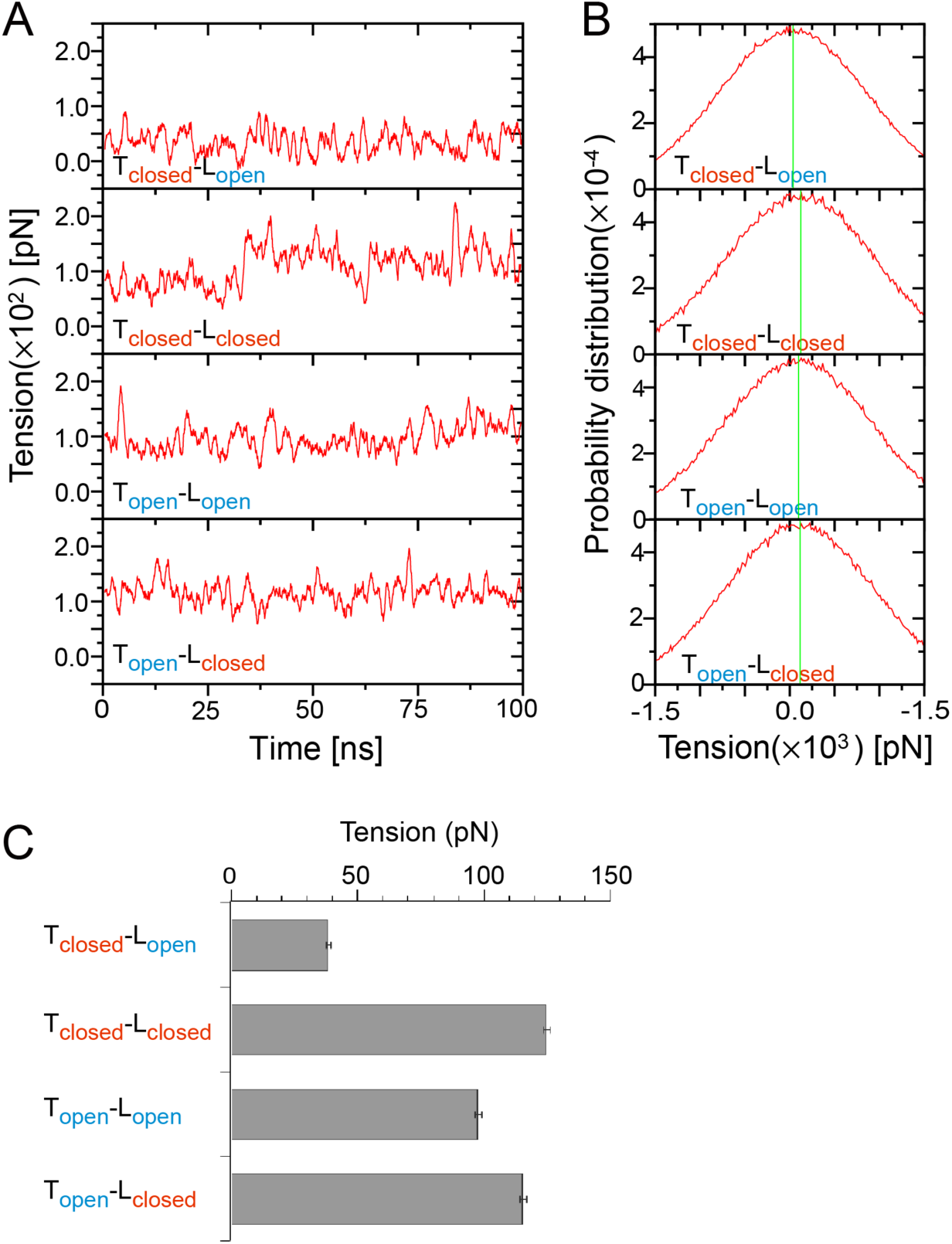
Estimation of tension in the disordered neck linker region using MD simulations of refined dimer models. **(A)** Time trajectories of the trends of tension during 100-ns simulations, using model systems (Fig. S4 D) derived from refined dimer models with various open and closed head combinations (Fig. 6). Tensions were calculated by multiplying the deviation of the Cα atoms at both ends of the disordered neck linkers from the potential minimum. The trend at each time *t* was calculated by averaging raw tension values within the time range [*t*-0.5: *t*+0.5] ns. Trends converged at ∼35 ns for the T_closed_-L_closed_ model and within 2 ns for the other three models. **(B)** Distributions of tension in disordered neck linkers, obtained from MD simulations. Green vertical lines indicate time-averaged tension for each state after convergence. **(C)** Time-average tensions and statistical errors for each state: 38.7 ± 0.8 pN for T_closed_-L_open_, 124.9 ± 1.5 pN for T_closed_-L_closed_, 98.1 ± 1.2 pN for T_open_-L_open_, 115.7 ± 1.2 pN for T_open_-L_closed_.

### Effect of tension of the neck linker on the tethered head binding

The role of tension applied to the neck linker on processive motility of kinesin dimer has been explored through the use of mutant kinesins with artificially extended neck linkers. Past research found that the velocity of the mutant’s processive motion decreased, but the ATP turnover rate remained unchanged compared to the wild-type (Yildiz et al., 2008; Shastry and Hancock, 2010; Clancy et al., 2011; Andreasson et al., 2015). These findings are consistent with the above hypothesis, as it predicts that reduced tension on the neck linker would facilitate the transition to the Topen-Lopen state, and additional ATP consumption is needed to return to the one-head-bound state, impairing the coupling between ATP hydrolysis and the forward step. To gather more direct evidence for this hypothesis, we systematically examined the relationship between the amount of tension applied to the neck linker and the probability of transitioning to two-head-bound states in the absence of ATP.

We used neck linker-extended mutants with varying levels of tension and observed their impact on the transition to the Topen-Lopen state. To increase the contour length of the disordered neck linker, we inserted poly-Gly residues of different lengths between the last two junctional residues of the neck linker (L335 and T336 residues; Fig. 8 A) (Yildiz et al. 2008; Clancy et al., 2011; Isojima et al., 2016). Then, we used a single-molecule FRET (smFRET) sensor to detect the transition between one-head-bound and two-head-bound states of the neck-linker extended mutants. Cysteine residues were introduced at the tip of one head (215Cys) and the base of another head (43Cys) on the cysteine-light background and were labeled with donor and acceptor dyes (Fig. 8 A). Previous studies showed that in the presence of a non-hydrolyzable ATP analogue, kinesin with this smFRET sensor displayed low (10%) and high (90%) FRET efficiencies, representing the two-head-bound state (Mori et al., 2007). In contrast, with a low amount of ADP, kinesin exhibited a distinct median (∼30%) FRET efficiency, indicating a one-head-bound state. To occasionally revert the transition towards the one-head-bound state and observe the ATP-independent transitions from one-head-bound to two-head-bound state, the observations were carried out in the presence of 2 µM ADP (ADP binds a few times per second, assuming ADP binding rate of 2 µM^-1^s^-1^ (Cross, 2004)).

**Figure 8.**
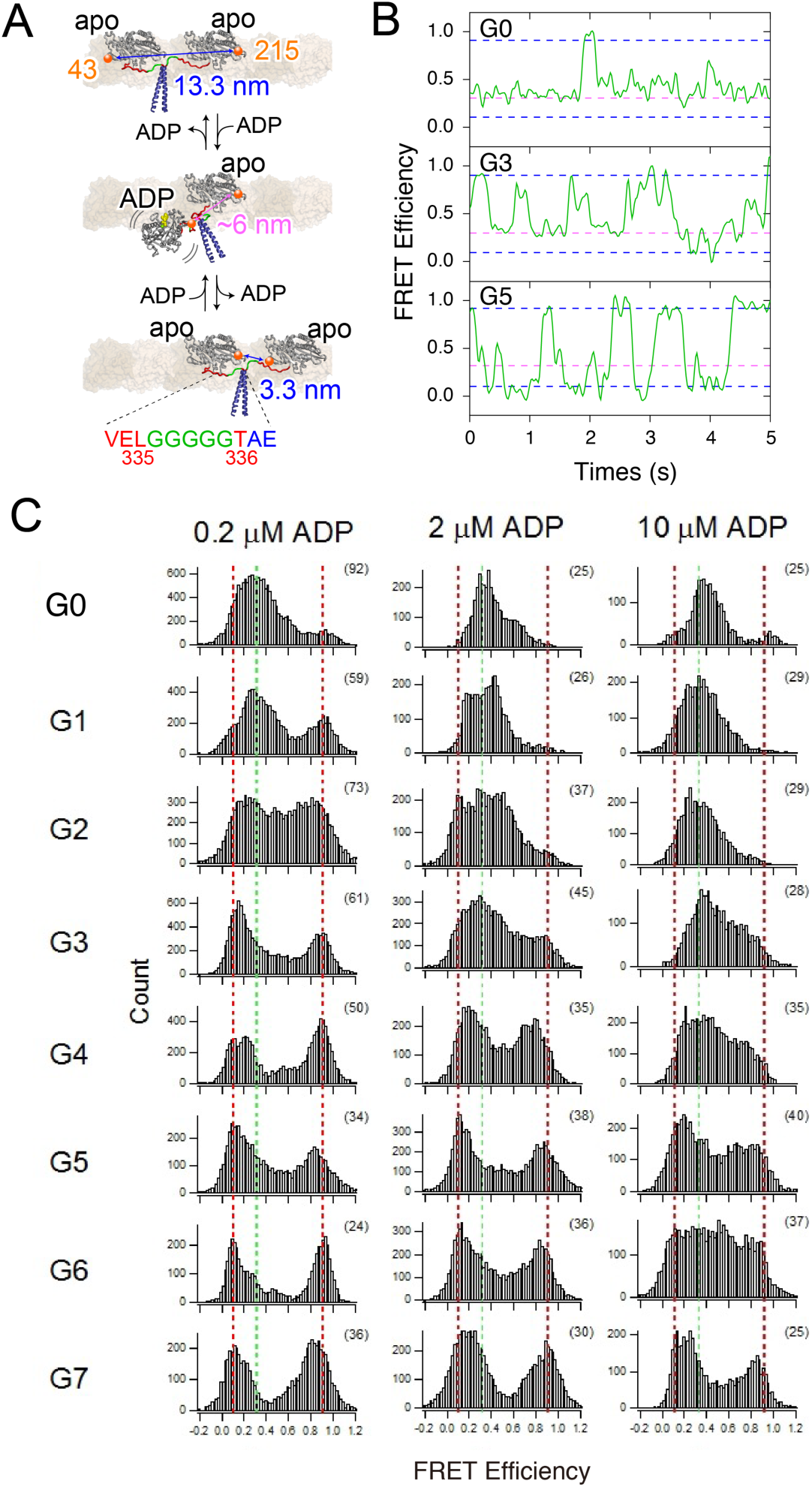
Single-molecule FRET observations of conformational transitions of kinesin dimer with reduced neck linker tension. (**A**) Schematic depicting off-pathway transitions from the two-head-bound state to one-head-bound states of a neck-linker extended mutant (5 poly-Gly insertion; green) without ATP. Positions of cysteine residues for labeling with FRET probe (215Cys on one head and 43Cys on the other head) are shown in orange spheres. (**B**) Typical FRET efficiency traces of Cy3/Cy5-labeled wild-type (G0) and neck linker extended mutants (3- and 5-Gly insertions; G3 and G5) bound to axonemes in the presence of 2 µM ADP. Blue and red dotted lines show previously reported peak FRET efficiencies for two-head-bound and one-head-bound states respectively (Mori et al., 2007). (**C**) Histograms of FRET efficiencies (from each frame of images) of Cy3/Cy5-labeled kinesin with varied neck linker lengths bound to axonemes in the presence of 0.2, 2 or 10 µM ADP. Occasional ADP binding to kinesin in the two-head-bound state reverts to one-head-bound state, and this frequency increases with higher ADP concentrations. Parentheses show the numbers of molecules analyzed. Dotted lines highlight peaks characteristic of putative two-head-bound (blue) and one-head-bound (red) states.

SmFRET observations revealed that the kinesin mutant, which has its neck linkers extended by inserting 3 poly-Gly residues (referred to as G3), frequently transitioned to two-head-bound states and then reverted back to a one-head-bound state triggered by occasional ADP-binding (Fig. 8 B). Under the same condition, the wild-type kinesin, without neck linker extension (termed G0), rarely showed transitions to the two-head-bound state. When the neck linker was further extended by 5 poly-Gly insertions, the duration of the one-head-bound state became negligible (Fig. 8 B), resulting in the mutant spending most of its time in two-head-bound states. The distribution of FRET efficiency shows that the G5 mutant occupies both high and low FRET states (i.e., two-head-bound states where either the donor- or acceptor-labeled head is leading) with nearly equal probability (Fig. 8 C), indicating that the detached head can bind to both forward and rearward tubulin-binding sites. The population of the medium FRET state (one-head-bound state) decreased as the number of inserted residues increased, and the two-head-bound states dominated for mutants with more than 5 poly-Gly insertions (Fig. 8 C). We reasoned that the G5 mutant could transition quickly from one-head-bound to Topen-Lopen states due to a reduced tension applied to the neck linker compared to the wild-type (98 pN). We used MD simulation to estimate this tension in the G5 mutant’s Topen-Lopen state and found that the tension was reduced to 19 pN (Fig. S5), which is now comparable to that in the Tclosed-Lopen state of wild-type (39 pN). These findings support the theory that the amount of tension applied to the neck linker is critical in regulating the transition from one-head-bound to two-head-bound states.

## Discussion

In the present study, we investigate the role of the neck linker in preventing the premature binding of the unbound head to the microtubule before ATP-binding to the leading head triggers the forward stepping of the head. High-resolution structural observations and the principal component analysis of kinesin structures revealed that the nucleotide-free structures belong to the distinct “open” conformational state (Fig. 3 A); in this state, the bulge of the C-terminal end of α4 on the surface of the head is located just forward from the C-terminus end of α6, to which the neck linker is attached (Fig. 4 D). This creates a steric hindrance when the neck linker is pulled forward but does not interfere when it is pulled backward (Fig. 6 A). MD simulation demonstrated that the tension applied to the neck linker in the Topen-Lopen state is over twice as high as that in the Tclose-Lopen state (Fig. 7 C). While the steric hindrance is small (approximately a 0.8 nm hemisphere), the tension increases significantly when the extension occurs during a rapidly increasing phase of the nonlinear force-extension curve of the entropic spring (Hariharan and Hancock, 2009). SmFRET observations of neck linker extended mutants further support the notion that the transition from one-head-bound to two-head-bound Topen-Lopen state is prohibited due to an intolerable tension increase (Fig. 8).

This asymmetric constraint on the neck linker is relieved when ATP binds and the kinesin motor domain transitions from open to closed state. During this transition, both the Front- and Rear-domains rotate anticlockwise relative to the Bottom-domain (Cao et al., 2014; Benoit et al., 2023; Fig. 4 B; Video 1). The R-domain, in particular, showcases a substantial rotational movement that leads to the closure of the nucleotide pocket on the left side and translates the distal end of α6 along with β1 away from α4 on the right side (Sindelar, 2011). As a result, the distal end of the α6 helix has room to extend, and K323 and T324 residues, previously excluded from the core in the nucleotide-free state, now contribute to an additional half-turn of α6 (Figs. 1D and 4D; Video 2). Simultaneously, the rotation of the F-domain unveils a hydrophobic pocket between α4 and β1. The formation of the α-helix by K323 and T324 moves the neighboring I325 residue close to the hydrophobic pocket and reorients its hydrophobic side chain, allowing the Ile side chain to bind complementarily to the hydrophobic pocket—an essential step in initiating neck linker docking (Vale and Milligan, 2000; Cao et al., 2014, Benoit et al., 2023). Finally, the rotation of the F-domain exposes the groove where the remainder of the neck linker forms two β-sheets and docks onto the head, completing the neck linker docking.

Because the tension applied to the disordered neck linker is only effective during the two-head-bound state, previous studies have mainly focused on how the tension of the neck linker influences the transition from a two-head-bound to a one-head-bound state (Block, 2007; Valentine and Gilbert, 2007; Gennerich and Vale, 2009). Here we show that the tension applied to the neck linker is also essential in regulating the transition from the one-head-bound to the two-head-bound state (Fig. 9, red rectangle). Since the conformational state doesn’t strictly correlate with the nucleotide state, in the following section, we’ll represent each state during the cycle as the nucleotide state/conformational state. The tethered ADP-bound head is in the semi-open conformational state and might transiently bind to the forward or rearward tubulin-binding site relative to the microtubule-bound nucleotide-free/open head. We did not estimate the tension in this two-head-bound state because the high-resolution structure of a semi-open state bound to the microtubule is not available as this state is unstable and transient (Benoit et al., 2021). Therefore, we can only speculate that the tension would lie somewhere between that of the Tclose-Lopen and Topen-Lopen states, and that microtubule binding of the tethered semi-open head may be restricted because the disordered neck linkers would need to adopt highly stretched configurations. Even when the tethered head managed to bind to the microtubule, the conformational transition of the ADP-bound head from semi-open to open state is prohibited as it would associate a significant increase in the tension on the neck linker, thereby maintaining the one-head-bound state. ATP-binding to the microtubule-bound head then induces rotational movement of the R-domain, moving the C-terminal of α6 upward from the bulge of α4, diminishing the asymmetric constraint on the neck linker. Under this condition, the tethered head binds to the forward tubulin-binding site and can transit to an ADP-bound/open state because the neck linker tension in this state is tolerable, followed by rapid ADP release from the head. Here, we highlight that the neck linker tension influences the allosteric transitions among three conformational states, not the nucleotide state directly.

**Figure 9.**
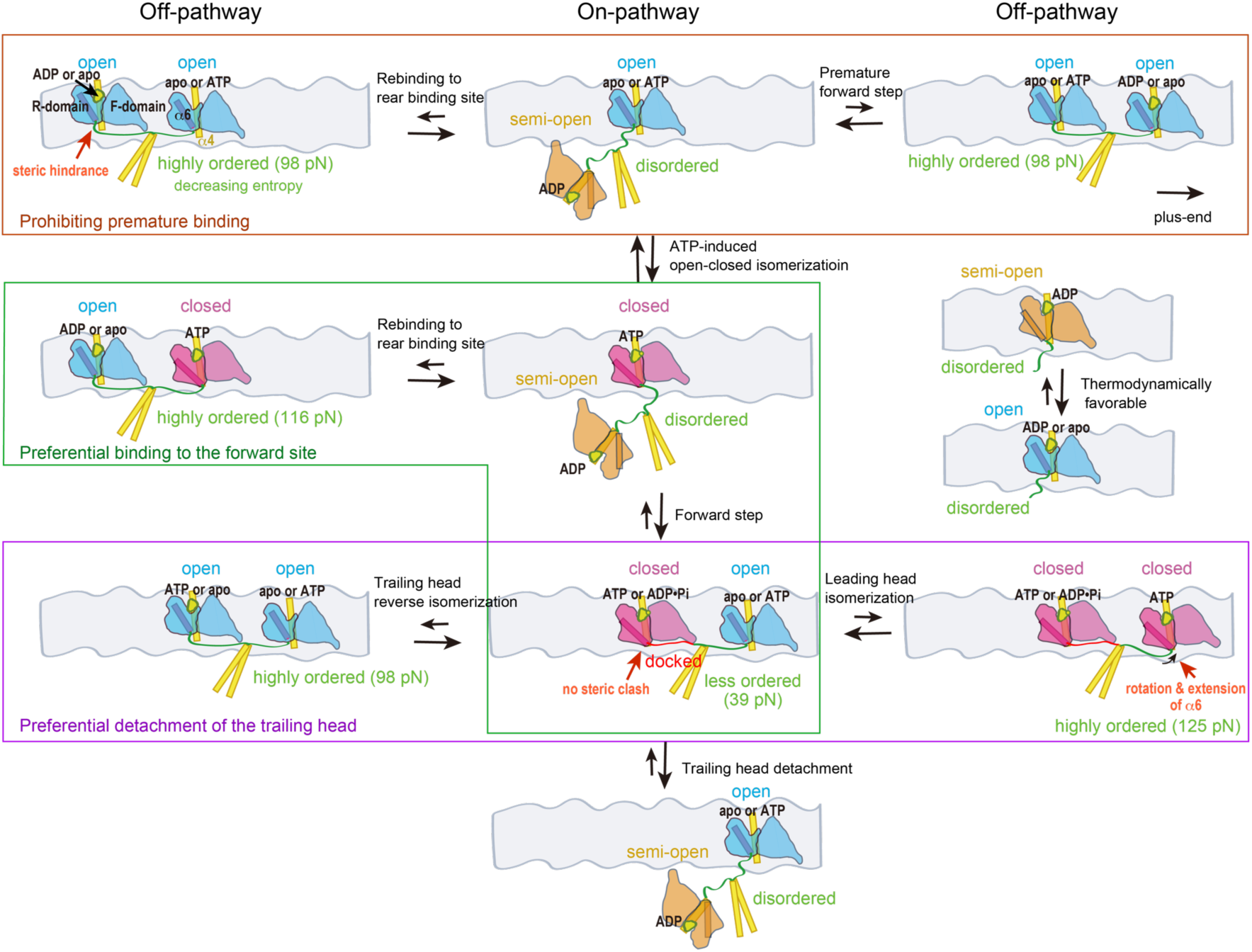
Tension-based regulation mechanism explaining the coordinated stepping of kinesin-1 along the microtubule. This model proposes that thermodynamically favorable transitions between a head’s conformational states (semi-open to open, and open to closed) are regulated by entropy loss due to neck linker stretching in dimeric kinesin. The semi-open, open and closed conformational states of the head are indicated in orange, blue and red, respectively, with the α6 helix, which connects to the neck linker, highlighted as a rod. The neck linker is shown in green, while the α4 helix, which directly interacts with the microtubule, and the neck coiled-coil are yellow. Premature binding of the tethered semi-open head to the microtubule in the ATP-waiting state is prevented by an intolerable increase in neck linker tension, as it causes substantial entropy reduction (red rectangle). This contrasts with a monomeric head with a disordered neck linker, where the semi-open-to-open transition is thermodynamically favorable (right). Even after ATP-induced isomerization of the microtubule-bound head, increasing tension prohibits rebinding of the tethered head to the rear-binding site (green hexagonal box), enabling preferential binding to the forward site. In the two-head-bound state, the trailing and leading heads stabilize in closed and open conformations, respectively, explaining why the trailing head hydrolyzes ATP and detaches from the microtubule before the leading head (purple rectangle).

The transition from semi-open to open conformation associated with ADP release is thermodynamically favorable for a head without neck linker constraint upon encountering the microtubule (Cross, 2004; Atherton et al., 2014; Benoit et al., 2023). However, in the case of dimeric kinesin during processive movement, this transition is also influenced by the level of neck linker tension; if the entropy reduction caused by stretching the disordered neck linker region exceeds the free energy decrease associated with the semi-open to open conformational transition of the head upon microtubule binding, the transition would become thermodynamically unfavorable (i.e., the sum of the free energy change of the head and the disordered neck linker increases). When the neck linker tension is reduced by inserting more than 5 poly-Gly residues, the loss in entropy decreases, making the transition to the Topen-Lopen state favorable.

This tension-based regulation mechanism (suppression of the thermodynamically favorable conformational transition by decreasing entropy) can be applied to other allosteric conformational transitions that alter the extensibility of the neck linker. For instance, when ATP binds to the leading head of the Tclose-Lopen state (Fig. 9, purple rectangle), the transition of the leading head from open to closed conformation (Tclose-Lclose state) increases the neck linker tension from 39 pN to 125 pN, under the condition that only the initial segment (residues 323-325) of the neck linker is docked onto the head (Figs. 6 B and 7). We included the I325 residue as it is crucial to thermodynamically stabilize the hydrophobic pocket exposed to the solvent in the closed state (Vale and Milligan, 2000; Sindelar, 2011, Cao et al, 2014). This excessive increase in tension would even prevent the initial I325 residue from docking onto the head. Therefore, even after ATP binding, the closed state is unstable in the leading head, and the head remains in the ATP-bound/open state (Benoit et al., 2023), which may follow by reversible ATP release. This sharply contrasts with the trailing head in the Tclose-Lopen state, which is stabilized in the ATP-bound/closed state due to the forward strain applied to the neck linker, because the reverse transition of the trailing head from the ATP-bound/closed state to the ATP-bound/open state (from Tclose--Lopen to Topen-Lopen state) is restricted by tension increase, as previously explained. ATP-hydrolysis and Pi release would then proceed in the stabilized closed nucleotide-pocket (Parke et al., 2010). Our kinetic measurements, which distinguished between leading and trailing heads, revealed that these heads are stabilized in low and high ATP-affinity state, respectively, and the neck linker strain controls the equilibrium between these states (Niitani et al., 2024). These findings further support the idea that neck linker tension influences the allosteric conformational transition, rather than directly affecting the nucleotide state. Hence, the tension-based regulation mechanism provides a rationale for the alternating catalysis of two heads during processive movement.

Moreover, this tension-based regulation mechanism provides a distinct mechanism from the neck linker docking model described earlier to explain the preferential forward stepping of the unbound head (Fig. 9, green hexagonal box). Assuming the microtubule-bound head binds ATP and transitioned to a closed state—while its neck linker remains detached—the tethered head can freely diffuse to both forward and rearward tubulin-binding sites. This assumption stems from the discovery of a relatively small free energy change associated with neck linker docking (Rice et al., 2003). However, the transition from semi-open to open state happens only at the forward binding site. We created a model of the Topen-Lclose dimer (after the tethered head binds to the rear binding site) where the leading head is in the closed state, but its neck linker is mostly detached from the head, leaving the first two amino acids to form an additional α6 helix (Fig. 6 C). The estimated tension in this state, according to an MD simulation, was 116 pN, which is even higher than that in the Topen-Lopen state (Fig. 7). Thus, this transition is thermodynamically unfavorable, and the transition to the Tclose-Lopen state will likely occur instead. In this model, neck linker docking is not necessary to account for the differences in neck linker tension (or entropy reduction); instead, it relies on the presence or absence of steric hindrance near the base of the neck linker in the trailing head. This idea is consistent with the findings of Taniguchi et al. (2005), who observed temperature-dependent stepping under load and demonstrated that the directionality of the kinesin step primarily originated from an entropic asymmetry. The two mechanisms explaining preferential forward stepping are not mutually exclusive. However, the tension-based mechanism may be crucial in preventing the tethered head from rebinding to the rear binding site, even after ATP-induced isomerization, especially under hindering load. Future studies are required to experimentally verify these models and gain a better understanding of how neck linker tension controls chemical and mechanical transitions of kinesin-1 during hand-over-hand motion. Additionally, studies are needed to examine whether this mechanism extends to other kinesin subfamilies with different neck linker properties, such as varying neck linker lengths (kinesin-2: Hariharan and Hancock, 2009; kinesin-3: Benoit et al., 2024) or unique interactions with the motor domain (kinesin-6: Guan et al., 2016; Ranaivoson et al., 2023).

## Materials and Methods

### DNA cloning and protein purification

Cysteines (S43C, E215C, T328C) or mutation in the switch II loop (G234A or G234V) were introduced into a “cysteine-light” human ubiquitous kinesin-1 349-residure monomer with C-terminal His6 tag (K349CLM), 336-residure monomer with N-terminal His6 tag (K336CLM), or 490-residue heterodimer with C-terminal His6 tag and Strep-tag (Rice et al., 1999; Tomishige et al., 2006; Mori et al., 2007). For the neck linker mutations, poly-glycine residues were inserted between L335 and T336 residues (Isojima et al., 2016), by PCR cloning.

Monomers and heterodimers were purified as described previously (Tomishige et al, 2000; Tomishige et al., 2006). For crystallization of nucleotide-free kinesin, K336CLM with G234A or G234V mutation was purified as described (Tomishige and Vale, 2000) and the elution from Ni-NTA resin was diluted in the buffer containing 20% (w/v) sucrose, 1 mM DTT, and 50 U/ml apyrase (Sigma-Aldrich). Then, the purified kinesin supplemented with apyrase was dialyzed against 25 mM PIPES (pH 7.0), 100 mM NaCl, 1 mM EGTA, 2 mM MgCl2 and 1 mM DTT for 1-2 hours to remove nucleotides in the solution, and further incubated with apyrase for 12-16 hours at 4°C to hydrolyze residual ATP and ADP. The sample was then purified using MonoQ column as described (Case et al, 1997; Tomishige and Vale, 2000), except that buffers without ATP were used and the column was equilibrated with 25 mM PIPES (pH 6.8), 1 mM EGTA, 2 mM MgCl2, 1 mM DTT and 50 mM NaCl. For crystallization of ADP-bound kinesin, K349CLM with G234A mutation was used, which was purified as describe above except that dialysis and incubation with apyrase were omitted. For cryo-EM observation, K349CLM was used.

### Crystallization, data collection and structural determination

Purified proteins with apyrase treatment were concentrated to 3-4 mg/ml in a crystallization buffer (10 mM Mops-NaOH (pH 7.0), 1 mM EGTA, 1 mM MgCl2, 1 mM DTT, 50 mM NaCl, and 20 % (w/v) sucrose). Crystallization was carried out at 4°C by the sitting drop vapor diffusion method with a subsequent streak-seeding. Initial crystals were obtained using a G234V mutant kinesin, which was purified with the same procedures with the G234A mutant, under a reservoir solution conditions of 20-21% PEG3350, 100 mM HEPES-NaOH, pH 7.7-8.0, 200 mM ammonium acetate, 3% xylitol (PS1). G234A mutant crystals were grown in the PS1 by streak-seeding method using a crystal of G234V mutant as a crystal seed. Then, G234A crystal for diffraction was obtained by additional streak-seeding using the crystals of G234A as a crystal nuclei, to exclude the contamination of G234V protein in the crystal. For ADP-bound G234A, protein was concentrated to ∼15 mg/ml in the crystallization buffer. Crystals were obtained under reservoir solution conditions of 23% PEG3350, 100 mM Bis-Tris pH 5.5 at 20°C. The crystals were transferred into each reservoir solution supplemented with 24% glycerol as a cryoprotectant and flash-cooled at 95K. X-ray diffraction data of apo and ADP-bound kinesin were collected on the BL-5A and NW-12 beamlines (wavelength l = 1.00 Å) at the Photon Factory, respectively. All X-ray diffraction data were integrated and scaled using the program XDS (Kabsch, 1993). The unit cell parameters are described in Table 1.

The structures were determined by the molecular replacement method using the coordinates of human kinesin-1 ADP-like structure (PDB# 1BG2; Kull et al., 1996) by the program Phaser (McCoy et al., 2007). Each initial model was refined and rebuilt using programs CNS (Brünger et al., 1998) and Coot (Emsley et al., 2010). In the final refinement process, the program Refmac5 (Murshudov et al., 1997) was used. The data collection and refinement statistics are summarized in Table 1. We expect that the relatively high R-factor for nucleotide-free G234A (Table 1) is likely due to the weak electron density (higher disorder) for the α0 and β2a region.

### Cryo-EM sample preparation, image analysis and model building

For high resolution cryo-EM observation, frozen grids of kinesin-decorated microtubules were prepared and imaged as described (Nishida et al., 2020) with some modifications. GMPCPP-microtubule (0.03 mg/ml polymerized in BRB80 (80 mM PIPES, 1 mM EGTA, 2 mM MgSO4)) was decorated with monomer kinesin K349CLM described above (0.2 mg/ml) in BRB80 on glow-discharged, carbon-coated grid (Quantifoil R 1.2/1.3, Cu/Rh, 300 mesh) supplemented apyrase (5 U/ml). Blotting and vitrifying were done using Vitrobot (Thermo Fisher Scientific). Cryo-EM data were acquired on a 200-KeV Talos Arctica electron microscope (Thermo Fisher Scientific). Data collection was done as described (Nishida et al., 2020). A total of 2,224 movie stacks were recorded. Image of helical 14-protofilament microtubules were processed with Relion-3 (Zivanov et al., 2018), PyFilamentPicker (Nishida et al., 2020) and Frealign (Grigorieff, 2007), as described (Nishida et al., 2020), except that the CTF parameters were estimated by CTFFIND 4 (Rohou and Grigorieff, 2015). After 3 cycles of pseudo-helical refinements, 74,859 particles (8 nm repeat of microtubule) were selected for several cycles of 3D auto-refinement followed by CTF refinement and Bayesian polishing in Relion-3. In total, data representing ∼1,050,000 asymmetric units from 2,438 microtubules were averaged. The overall gold-standard resolution was 3.48 Å. The local resolution was calculated with MonoRes (Vilas et al., 2018) and the output resolution map was used for local sharpening with LocalDeblur (Ramírez-Aportela et al., 2020).

To develop an atomic model for kinesin-microtubule complex, the atomic structure of tubulin (PDB# 3JAT; Zhang et al., 2015) and our G234A apo crystal structure were fit in the sharpened map using UCSF chimera (Pettersen et al., 2004). For model refinement, we used the MDFF package (Trabuco et al., 2008) as described (Nishida et al., 2020), and PHENIX (Adams et al., 2010). Refined model was used for further sharpening of maps with LocScale (Jakobi et al., 2017).

### Cryo-EM sample preparation, image analysis for gold-labeled kinesin

For gold-cluster labeling, purified K349CLM with T328C mutation was labeled with monomaleimide-undecagold as described (Kikkawa et al., 2000). Free gold-cluster and unlabeled kinesin were removed through RESOURCE S column (1 ml, GE Healthcare) chromatography. The efficiency of labeling was ∼80% as estimated using SDS-PAGE. The eluted fraction of the gold-labeled protein was concentrated with Microcon-YM-3 (Millipore), and then dialyzed against a BRB12 solution (12 mM PIPES (pH6.8), 1 mM EGTA and 2 mM MgCl2) using micro-dialysis device with 10k MWCO dialysis membrane (Spectrum Laboratories) for 1 hr immediately before the EM preparation.

Frozen grids of gold-labeled kinesin-decorated microtubules were prepared and imaged as described (Kikkawa et al., 2000) with some modifications. Microtubule (0.8 mg/ml) was decorated with gold-labeled kinesin heads (0.6 mg/ml) in BRB12 buffer supplemented 20 mM taxol and apyrase (5-10 U/ml). Cryo-EM images were recorded with a JEM-3000SFF electron microscope (JEOL Ltd., Tokyo, Japan) equipped with a field emission gun and a super-fluid helium stage (Fujiyoshi et al., 1991) and operated at 300 kV. Images were recorded on SO-163 (Kodak, USA) at a nominal magnification of 40,000x, using a 2-second exposure and a total electron dose of 20 electrons / Å^2^. The micrographs were developed for 14 min at 20°C using full-strength D19 developer. Selected films were digitized at a step size of 10 μm with a CCD film scanner LeafScan 45 (Scitex), which corresponds to 2.5 Å on the samples.

Cryo-EM images were processed with a script system, Ruby-Helix (Metlagel et al., 2007). Images of helical 15-protofilament microtubules were analyzed as described. In total, data representing ∼9,000 asymmetric units in 33 segments from four microtubules were averaged.

### Principal component analysis (PCA) of kinesin structures

PCA of kinesin structures were performed on Bio3D package as described (Grant et al., 2007). Atomic coordinates for kinesin superfamily crystal structures (247 motor domain chains extracted from 137 structures (until PDB# 6G6Z) deposited in PDB plus our G234A and G234V apo crystal structures and cryo-EM apo structure were used. Prior to the PCA, structures were superposed on the invariant “core” residues consisting of 31 residues (see Fig. 3 C, colored in gray; residues 10-15, 48-49, 80-85, 228-231, 297-303, 311-316, which are part of structural elements β1, β2, β3, β7, β8 and α6).

Subdomains that are invariant in the semi-open, open and closed states were identified with the program GeoStas (Romanowska et al., 2012), using PDB# 5LT1 (Cao et al., 2017) as semi-open state, our EM-apo structure as open and PDB# 4HNA as closed states.

### Modeling of kinesin-tubulin complex for semi-open and closed states

Kinesin-tubulin complex structures were constructed by fitting crystal structures on the cryo-EM maps by UCSF Chimera. The crystal structures of semi-open (PDB# 5TL1; Cao et al., 2017), and closed (#4HNA; Grant et al., 2013) kinesin-1 along with the tubulin dimer model from our cryo-EM-apo structure were fit to the cryo-EM density map of ADP-bound (EMD-5164) and ADP-AlFx bound (EMD-6188) kinesin-tubulin complex (Sindelar and Downing, 2010; Shang et al., 2014).

### Modeling of dimeric kinesin and refinement using molecular dynamic simulation

For dimeric kinesin models, two kinesin head-tubulin complexes created as described above were fitted to adjacent tubulin-dimer portions within single protofilament of the cryo-EM density map (EMD-1340; Sindelar and Downing, 2007). The coordinates for the disordered neck linker region and neck coiled-coil were modeled based on those of dimeric kinesin (PDB#3KIN; Kozielski et al., 1997) and loop modeling by satisfaction of spatial restraints were performed using MODELLER program (Fiser et al., 2000). Then MD simulations were applied to the manually modeled neck linker to refine the positions of the disordered neck linker region. The MD calculation refines the neck linker configuration but does not significantly affect the end-to-end distance of the neck linker.

The MD simulations of the disordered neck linker were performed by GROMACS 4.0.2 (Berendsen et al., 1995; Lindahl et al., 2001) with the AMBER-99SB for proteins (Lindahl et al., 2001; Sorin and Pande, 2005; DePaul et al., 2010) and the TIP3P for waters. The simulation system contained the disordered neck linker, neck coiled-coil, all the atoms of the kinesin motor domain and tubulin within 12 Å from the neck linker, and a few additional residues for continuity, which was solvated with waters and counter ions in a box with water layers at least 9 Å from the protein. The position restraints were applied to all the non-H atoms except the disordered neck linker and the neck coiled-coil, and to the main chain atoms at the boundary residues between the head and the neck-linker; K323 residue in the leading head and E334 (Tclosed) or K323 residue (Topen) in the trailing head. To inhibit unzipping of the coiled-coil, we added a harmonic potential between the Cα atoms of the residue 337 in both chains. All the systems were first energy-minimized and waters and ions were then equilibrated for 1 ns. The production runs were performed for 30 ns for Tclosed-Lopen, 20 ns for Tclosed-Lclosed, 16 ns for Topen-Lopen, and 30 ns for Topen-Lclosed, in which the coordinates were saved every 2ps (every 1ps for the Topen-Lopen state). The simulations employed the NPT ensemble (Parrinello and Rahman, 1981; Berendsen et al., 1984) with 300 K and 1 bar, the periodic boundary conditions, the particle mesh Ewald (Darden et al., 1993) with cut-off 1.0 nm for the electrostatics, the van der Waals cutoff 1.4 nm, the LINCS for constraining bond lengths (Hess et al., 1997), and 1 fs and 2 fs time step in the production run for the Topen-Lopen and for the others, respectively. For each simulation, convergence was assessed by a few measures.

### Estimation of the tension applied to the neck linker

The MD simulations for the estimation of tension posed to the disordered neck linker in the refined dimer models were performed by GROMACS 4.5.5 with the AMBER-99SB for proteins and the TIP3P for waters. The simulation system contained the neck coiled-coil and the disordered neck linker region of which two ends are E334 for Tclosed or T328 residue for Topen in the trailing head and I325 residue in the leading head (Figure S4 D). These protein segments were solvated with waters and counter ions in a box with water layers at least 15 Å from the protein. The initial structures were taken from the refinement structures shown in Fig. 6. To evaluate the tension applied to the neck linker in each conditions, the distance constraints were applied to the Cα atoms of the edge residues in both ends of the neck-linker in the form of harmonic potential: k (R - R0)^2^, where R is the distance between Cα atoms on two ends, spring-constant k is set as 105 kJ/mol/nm^2^, constraint distance R0 is set as the corresponding average distance for each system during the refinement MD. To inhibit unwinding of the neck coiled-coil, we added a harmonic potential between the Cα atoms of the residue 337 in the two polypeptides as described above. Starting from the refined structures, all the systems were first energy-minimized and waters and ions were then equilibrated for 1ns. The production runs for estimation of the tension were performed for 200 ns for Tclosed-Lopen, and 100 ns for the other systems, in which the deviation of the distance from the constraint distance R0 were saved every 0.2 ps. The neck linker tension was calculated as the spring constant k multiplied by the deviation from the constraint distance R0. Other simulation protocols are the same as those described in the refinement-MD. For each system, convergence of the tension was confirmed (see Fig. 7 A).

### Single-molecule FRET observations

Single molecule FRET measurement and data analysis were carried out as described previously (Tomishige et al., 2006; Mori et al., 2007) using Cy3- and Cy5-labeled heterodimers (E215C in one chain and S43C in the other chain) with insertions of poly-Gly between L335 and T336 residues. Dual-labeled heterodimer molecules bound to the axoneme in the presence of 0.2-10 µM ADP and 5 U/ml hexokinase (which converts contaminating ATP to ADP) were excited with 514 nm argon laser (35LAP321; Melles Griot), and donor and acceptor fluorophores were simultaneously observed using a Dual-View (Optical Insights) and EM-CCD camera (iXon DV860; Andor) at an acquisition rate of 100 frames/s.

## Online supplemental material

Supplemental data and legends relating to Figs. 1, 2, 6, and 7 can be found in Figs S1, S2, S3, S4, and S5.

## Data availability

The data that support the findings of this study are available from the corresponding author upon request. The atomic coordinates and structure factors for the crystal structures have been deposited in the Protein Data Bank under the accession codes 9L6K (nucleotide-free G234V mutant), 9L78 (nucleotide-free G234A mutant), and 9L7E (ADP-bound G234A mutant). The cryo-EM density map has been deposited in the Electron Microscopy Data Bank under accession code EMD-62874, and the corresponding atomic coordinates have been deposited in the Protein Data Bank under accession code 9L7M.

## Acknowledgments

We thank M. Nakajima for assistance in the cloning of constructs, C. Sindelar for providing cryo-EM density maps, Y. Fujiyoshi for use of cryo-EM microscope for gold-labeled kinesin, R. Vale for helpful discussions. The synchrotron-radiation experiments were performed at the Photon Factory (2003S2-002 and 2008S2-001).

M. Tomishige, M. Kikkawa and T. Makino are supported by Research Grant for Young Investigators from Human Frontier Science Program. M. Tomishige is supported by Grants-in-Aid for Scientific Research on Priority Areas from MEXT, and by the Asahi Glass Foundation. M. Tanokura is supported by the National Project on Protein Structural and Functional Analyses and the Targeted Proteins Research Program of MEXT.

## Author contributions

M. Tomishige and T. Makino conceived and designed the experiments. T. Makino carried out purification, crystallization, structural determination and analysis with K. Miyazono and M. Tanokura. T. Makino and M. Kikkawa performed cryo-EM observation and analysis with Y. Komori and H. Yanagisawa. R. Kanada and S. Takada ran MD simulations. T. Mori and M. Tomishige carried out smFRET observations. T. Makino, R. Kanada and T. Mori prepared figures and M. Tomishige wrote the manuscript.

---

**Figure S1. Related to Fig. 1B.**
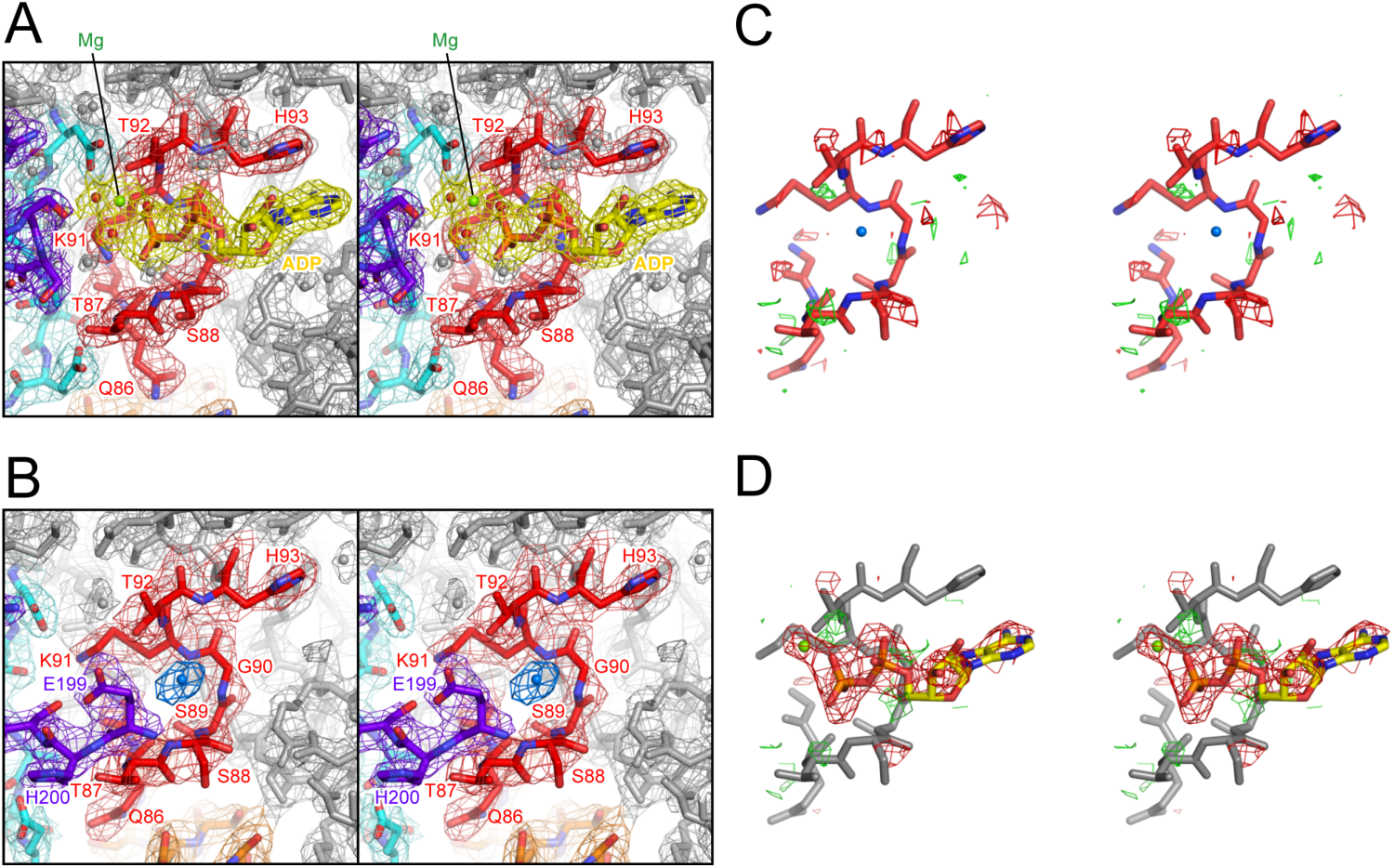
**(A,B)** Stereo-views of the 2*F*_o_-*F*_c_ maps (α = 1.0) around the nucleotide-binding P-loop (**A**: G234A kinesin complexed with ADP, **B**: G234A kinesin crystallized under nucleotide-free condition). ADP and Mg^2+^ ion are shown in yellow and lime green, respectively. **(C)** *F*_o_-*F*_c_ difference map of the nucleotide-binding P-loop of nucleotide-free G234A mutant. Positive (σ = 2.0) and negative (σ = -2.0) densities are shown in green and red, respectively. No residual density was observed around the putative water molecule (blue). **(D)** *F*_o_-*F*_c_ map (α = ±2.0) obtained by one round of refinement after replacing the putative water molecule with MgADP in the nucleotide-free G234A model. The negative density (red) further clarifies the absence of MgADP in the nucleotide-binding pocket.

**Figure S2.**
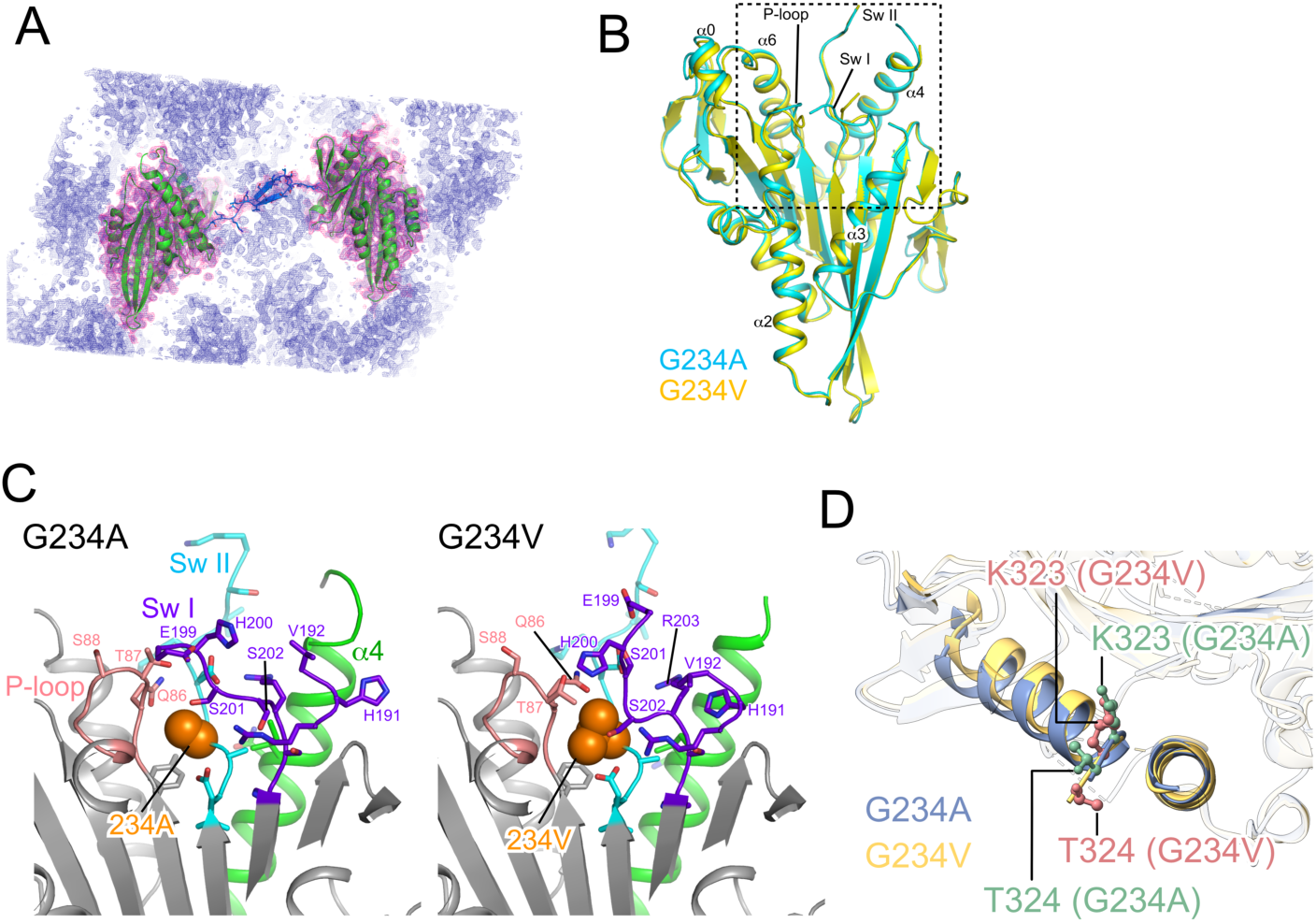
2.8 Å crystal structure of nucleotide-free G234V mutant kinesin. **(A)** The crystal packing of nucleotide-free G234V mutant. 2*F*_o_-*F*_c_ map is shown in mesh (dark-blue) and the densities around the asymmetric unit are colored in pink. The proximal ends of the neck linkers of chains A and B face each other, with a cylindrical cavity between them. This cavity allows an antiparallel β-sheet formation between the two stretched neck linkers of chain A and B (shown in blue). While this demonstrates the neck linker peptides have an ability to form antiparallel β-sheets, such an arrangement would be geometrically impossible for the two neck linkers in a dimer, as their C-termini are joined in parallel by the neck coiled-coil. In the G234A apo structure, we did not observe density corresponding to the antiparallel β-sheet in the cavity, likely due to its slightly smaller cavity size. **(B)** Comparison of the overall structures of nucleotide-free G234A (cyan) and G234V (yellow) crystal structures. G234V kinesin structure is nearly identical to that of G234A, except for the difference in the P-loop. **(C)** Close-up view of the nucleotide-binding pocket of G234A and G234V mutants (dotted rectangle in panel B). The side chains of residue 234 are shown as space-filling. The P-loop (pink) of G234V displayed a closed conformation, distinct from previously solved kinesin structures, indicating that the P-loop can adopt different configurations without bound phosphate. **(D)** Close-up view of the α4/α6 helices of nucleotide-free G234A (blue) and G234V (yellow) (the G234 structure is included from Fig. 1 C). The first two residues of the neck linker, K323 and T324, are highlighted as ball-and-stick models (G234A in green, G234V in red). In the G234V mutant, the T324 residue of the neck linker was visible and shifted toward the minus-end of microtubule compared to that of G234A.

**Figure S3. Related to Fig. 2.**
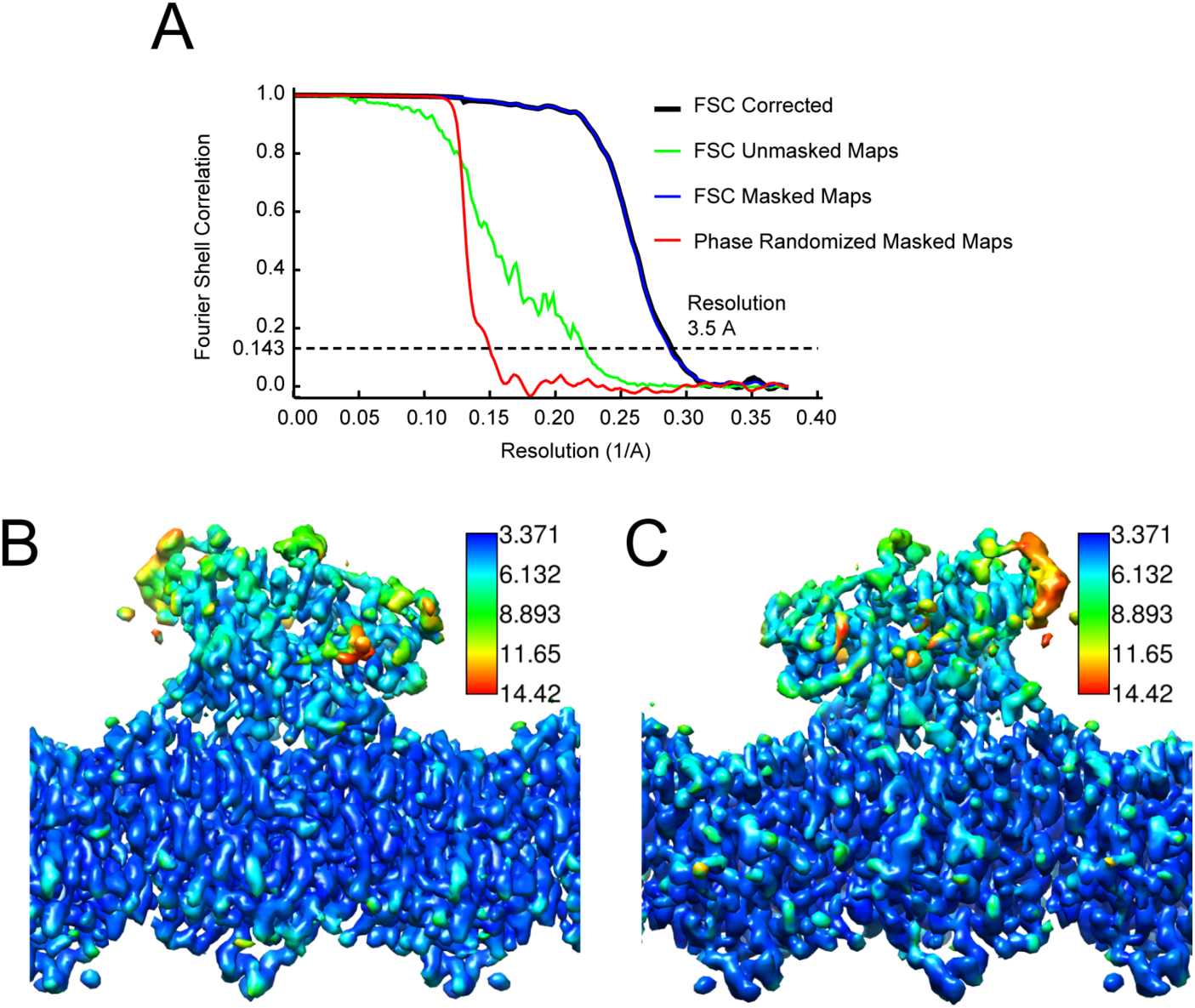
**(A)** Fourier shell correlation (FSC) curve for global resolution estimation, using a FSC 0.143 criterion. **(B and C)** Local resolution estimation from MonoRes (Vilas et al., 2018) with colors according to the color scale of left-side view (B) and right-side view (C). This local resolution map was used in local map sharpening (LocalDeblur (Ramírez-Aportela et al., 2020)).

**Figure S4. Related to Fig. 7.**
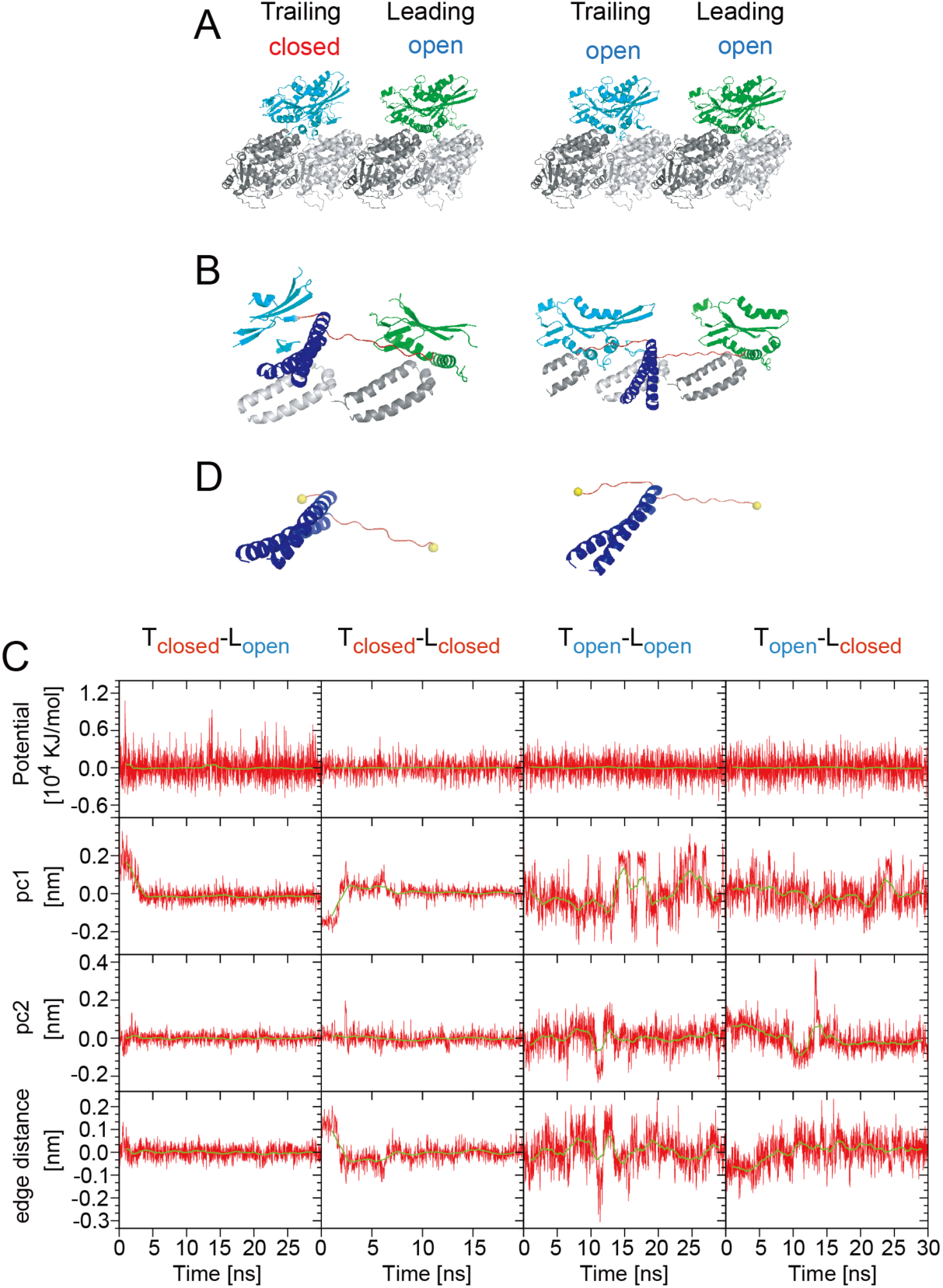
**(A)** Kinesin-microtubule complex models were initially created by fitting crystal structures to cryo-EM density maps (left: T_closed_-L_open_, right: T_open_-L_open_). Disordered regions—those invisible in the crystal structures—were excluded from this model. **(B)** The model systems used to estimate the thermally equilibrated conformation of the disordered neck linker region. Molecular dynamic (MD) simulations moved the disordered neck linkers (shown in red), while kinesin and tubulin residues within 12 Å from the neck linker (cyan, green) were position-constrained to act as steric hindrance. **(C)** Time trajectories of total potential energy, principal components (pc1 and pc2) from standard PCA, and end-to-end distances of the disordered neck linker, during MD calculation of four kinesin dimer models. End-to-end distances converged after 2-15 ns, indicating that the disordered neck linkers reached near-thermal equilibrium by the end of the simulations. Representative snapshots of equilibrium states shown in Fig. 6 were selected from structures after extended simulation periods. These structures had end-to-end distance and pc1/pc2 values nearly identical to the average values (< 0.01 nm deviation): *t* = 29.870 ns for T_closed_-L_open_, 19.920 ns for T_closed_-L_closed_, 12.758 ns for T_open_-L_open_, and 29.696 ns for T_open_-L_closed_. (**D**) Model systems used to estimate the tension on disordered neck linkers (Fig. 7). MD simulations were performed using the refined neck linker and neck coiled-coil (Fig. 6). Harmonic potentials were applied to the Cα atoms at both ends of the disordered neck linkers (yellow spheres; E334 for T_closed_ (left) and T328 for T_open_ (right), and I325 for L_open_). Tension was calculated by multiplying the deviation of the Cα atoms from the potential minimum by the spring constant. Time trajectories are shown in Fig. 7 A.

**Figure S5.**
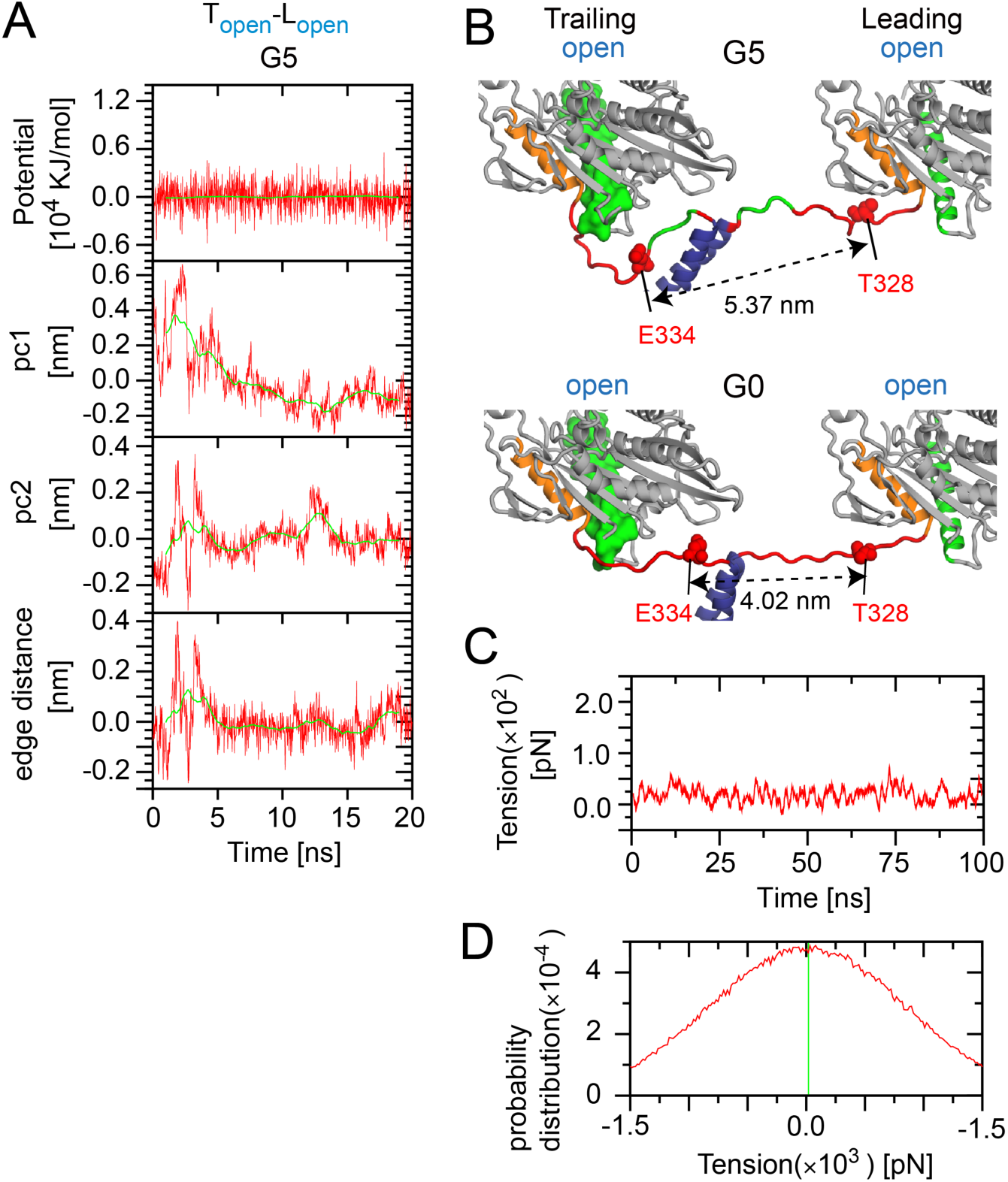
Refinement structure of disordered neck linkers and its estimated tension of neck linker-extended mutant in T_open_-L_open_ state. (**A**) Time trajectories of total potential energy, pc1 and pc2 from standard PCA, and end-to-end distance of the disordered neck linker during MD calculation of G5 mutant in T_open_-L_open_ model. (**B**) T_open_-L_open_ model of G5 mutant compared with the T_open_-L_open_ model without neck linker extension (G0). Thermally equilibrated conformation of the disordered neck linker was determined by MD simulation. Disordered neck linkers were isolated from the refined model, and harmonic potentials were applied to Cα atoms of residues at both ends (T328 in trailing head and I325 in leading head) to estimate tension. (**C**) Time trajectory of tension trends during 100-ns simulations. Trend at each time *t* was calculated by averaging raw tension values within the time range [*t*-0.5: *t*+0.5] ns. (**D**) Distribution of tension in disordered neck linkers from MD simulation. Green vertical line indicates time-averaged tension after convergence. Average tension, calculated using the data after the convergence, was 19.0 ± 1.2 pN.

Video 1. **Subdomain motions associated with the allosteric conformational changes of kinesin on microtubule (back-view).** The morph movie between semi-open, open and closed states, with subdomains color-coded. Related to Fig. 4 B.

Video 2. **Subdomain motions associated with the allosteric conformational changes of kinesin on microtubule (side-view).** The morph movie between semi-open, open and closed states, with subdomains color-coded. Related to Fig. 4 D.

Video 3. **Thermal motion of the disordered neck linkers in Topen-Lopen or Tapo-Lapo state.** The movie shows molecular dynamic simulation for 16 ns. The disordered neck linker regions are shown in stick representation. Related to Fig. 6 A (left).

Video 4. **Thermal motion of the disordered neck linkers in Tclosed-Lopen or TATP-Lapo state.** The movie shows molecular dynamic simulation for 30 ns. The disordered neck linker regions are shown in stick representation. Related to Fig. 6 A (right).

